# Enhanced Identifications and Quantification through Retention Time Down-Sampling in Fast-Cycling diagonal-PASEF Methods

**DOI:** 10.1101/2025.04.23.650190

**Authors:** Christopher R. Below, Oliver M. Bernhardt, Stephanie Kaspar-Schönefeld, Sander Willems, Edoardo Coronado, Ino D. Karemaker, Bettina Streckenbach, Luca Räss, Sandra Schär, Dennis Trede, Jonathan R. Krieger, Tejas Gandhi, Roland Bruderer, Lukas Reiter

## Abstract

Data-independent acquisition mass spectrometry is essential for comprehensive quantification of proteomes, enabling deeper insights into cellular processes and disease mechanisms. On the timsTOF platform, diagonal-PASEF acquisition methods have recently been introduced to directly and continuously cover the observed diagonal shape of the peptide precursor ion distributions. Although diagonal-PASEF has shown great promise, its broad adoption as a routine workflow has been hampered by a lack of available algorithmic solutions and a paucity of demonstrated real-world applications. Here, we conducted a systematic and comprehensive optimization of diagonal-PASEF for 17-minute gradients on the timsTOF HT in conjunction to Spectronaut 19. We demonstrate that Spectronaut 19 fully supports all tested diagonal-PASEF methods independent of the number of slices or overlaps and with minimal user intervention required. Using our optimized analysis strategy we coupled diagonal-PASEF acquisitions to retention time down-sampling by summation (RTsum), thereby providing a novel mode of analysis of this data that exploits the fast-cycling nature of diagonal-PASEF methods. Through the combination of RTsum with diagonal-PASEF, we demonstrate that this strategy yields higher signal-to-noise ratios while retaining the peak shape for analytes of interest. Importantly, combining RTsum with diagonal-PASEF improved overall identifications and quantitative precision when compared to dia-PASEF with static or variable quadrupole isolation widths and across different input amounts for cell line injections. We also tested the performance of diagonal-PASEF in controlled quantitative experiments where diagonal-PASEF outperformed dia-PASEF in the overall number of retained candidates, quantitative precision and identifications on peptide level. These data indicate that RTsum demonstrates a positive use case of the high sampling rate of diagonal-PASEF. Collectively, our findings imply that diagonal-PASEF methods are superior for peptide and protein level analyses especially at lower input amounts.

## Introduction

Proteomics studies the complete set of proteins encoded by a genome, and as such offers complementary information to the gene dosage governed by nucleic acids expression levels ^1^. The study of proteins provides mechanistic insights into signaling cascades or drug responses at cellular and even subcellular levels ^2^.

Historically, proteomics has lagged behind next-generation genome sequencing in terms of scale and depth due to the intricate workflows necessary to accurately capture the multifaceted biochemistry of proteins and peptides. The same biochemical complexity that enables proteins and peptides to drive diverse biology has also necessitated advances in sample preparation, analyte separation and data analysis, all of which presented significant challenges^3^. With the improvements in liquid chromatography mass spectrometry-based (LCMS) techniques, in combination with novel bioinformatics technologies including improved computing power and leveraging artificial intelligence, substantial decreases in data analysis time, as well as increases in proteome coverage have made high-throughput proteomics not only achievable but widely adopted^4,5^.

Contemporary LCMS approaches aim to yield high peptide and protein identifications while retaining quantitative accuracy and precision. Whereas data-dependent acquisition (DDA) used to be the gold standard for proteomics, data-independent acquisition (DIA) is increasingly replacing DDA especially for methods with high sample throughput. One of the main advantages of DIA is that it is not dependent on a precursor selection heuristic and thereby results in an unbiased sample survey and higher data completeness by fragmenting each precursor once within each DIA acquisition cycle^6,7^. In addition, precursors as well as corresponding fragment ions, provide elution profiles, thus improving the quality of fragment peaks and enabling MS2-level quantitation in addition to that obtained through consecutive MS1 scans. In recent years significant developments in instrumentation, data acquisition and data processing have considerably improved sensitivity and reproducibility of DIA workflows making it ideally suited for high-throughput large sample cohort studies ^8–10^. As a result of this, the boundaries of what is possible are continuously shifting in proteomics studies and this has been reflected in an exponential increase in sample throughput and downscaling of sample preparation to the single cell level^11,12^.

The advent of trapped ion mobility spectrometry (TIMS^13^) technology marked a significant advancement in proteomics, enhancing depth by increasing ion usage through parallel accumulation serial fragmentation (PASEF)^14,15^. TIMS introduces an additional dimension of separation, as well as molecular characterization via collisional cross-sections, thereby providing higher confidence and cleaner MS and MS/MS-spectra ^16^. Furthermore, TIMS increases selectivity through the spatial and temporal focusing of ions eluting from the ion mobility (IM) device and when applied in conjunction with PASEF allows for the near complete ion utilization throughout the analytical gradient. By combining DIA with TIMS, sample-complexity was shown to be further reduced in a scan mode introduced in 2020 termed dia-PASEF^16^. Within dia-PASEF acquisitions, predefined quadrupole isolation windows for ion selection in the m/z and mobility plane are applied in each TIMS elution. A variety of different window schemes have been introduced^17,18^, including the usage of variable window widths (py_diAid^19^), very narrow windows (thin-dia-PASEF^20^) and application of a method that continuously scans precursors with multiple horizontal quadrupole isolation windows (slice-PASEF^21^). In addition, such methods have also been used in conjunction with scan summation that has the promise to improve the signal-to-noise ratio for single cell-based analyses of fast-scanning timsTOF data^12^.

To further increase the analytical performance of DIA-based approaches on timsTOF mass spectrometers, researchers have recently proposed diagonal-PASEF acquisition methods termed either synchro-or midia-PASEF, which were designed to acquire the ion cloud more efficiently^22,23^. These methods operate by seamlessly and continuously following the observed diagonal shape of the precursor ion distribution in m/z and ion mobility dimension. The quadrupole isolation window is moved synchronously with the IM elution, which leads to better utilization of the trapped ion cloud and allows correlation of the fragment and precursor signal more tightly. The synchronized movement can slice the ion cloud into several, method dependent, TIMS ramps which are often termed diagonal-PASEF slices. Compared to conventional dia-PASEF approaches, the cycle time of diagonal-PASEF methods does not increase with an increased overall mass range as each scan covers the desired m/z range. This is particularly true for non-overlapping schemes, such as synchro-PASEF, where only a few slices are required to cover the range of interest, thereby resulting in very short cycle times and improved peak coverage as a result of more frequent sampling.

An initial study has shown that synchro-PASEF can achieve comparable proteome depth compared to classic dia-PASEF modes with superior quantitative performance as a result of the higher sampling rates, which clearly demonstrates the potential of diagonal-PASEF acquisition modes^18^. Software tools have already integrated initial data analysis algorithms for diagonal-PASEF (Spectronaut, DIA-NN, and alphaDIA), and while initial results show that such methods promise to cover the observed ion-cloud more efficiently, their utility is currently still hampered by a lack of systematic method evaluation and data analysis. Several parameters can be optimized to tailor the method for specific applications. Effective implementations could focus on maximizing sensitivity and minimizing cycle time for optimal peak coverage. Scan summation strategies have been described for dia-PASEF applications which hold the promise to improve the signal-to-noise ratio for single cell-based analyses of fast-scanning timsTOF data^12^. Unfortunately, to date, such approaches have not been applied to fast-scanning diagonal-PASEF methodologies. We therefore set out to develop a data analysis algorithm with optimized ion current extraction and scan summation for Spectronaut^6^. In addition, we optimized the diagonal-PASEF acquisitions on a timsTOF HT mass spectrometer for different applications and provided evidence for its benefit in modern proteomics applications.

## Results

### Spectronaut version 19 fully supports diagonal-PASEF acquisitions

To systematically test diagonal-PASEF LCMS methods we first had to extend the data analysis of Spectronaut to support this novel type of data. The main aspects of making Spectronaut support diagonal-PASEF was to change the data-extraction region that is used to extract MS2-features for any given MS1 event. In Spectronaut version 18, the predecessor of Spectronaut version 19, the MS2-features for any given precursor are only extracted from one dia-PASEF box that is defined as rectangle in 1/k0 – m/z space (**Figure 1, A**). As diagonal-PASEF conducts continuous quadrupole movements such rectangles are not applicable for these types of methods. We therefore adapted the data extraction algorithm to account for individual isolation windows per micro-scan within a diagonal-PASEF frame (**Figure 1, B**). To validate that the MS2-extraction worked sufficiently with the new data extraction rule in Spectronaut version 19, we investigated extracted-ion-mobilograms (XIMs) from precursors that were identified in a Hela acquisition using either a diagonal-PASEF MS-method with 4-slices or a dia-PASEF MS-method consisting of 12 PASEF-ramps. We investigated the protein HNRPR (Uniprot accession O43390) for which 48 unique precursors were identified in the diagonal-PASEF run and 51 were identified in the dia-PASEF acquisition (**Figure 1 C-D, Supplementary Figure 1, A**). Manual inspection of representative XIMs generated from MS/MS scans highlighted the absence of any artifacts or data-gaps especially between adjacent slices for the investigated diagonal-PASEF acquisition (**Figure 1 D**). In addition, the XIM of the selected precursor did not show any alterations between the dia-PASEF or diagonal-PASEF acquisition albeit a higher intensity signal was detected which confirmed previous findings from other studies^22^. We validated these findings in two additional precursors “_NLATTVTEEILEK_.2” which also mapped to HNRPR (**Supplementary Figure 1, B**) and “_DLLDLLVEAK_.2” which mapped to DDX3X (Uniprot accession O00571, **Supplementary Figure 1, C**). These data indicated that an efficient data-usage can be conducted using the upgraded data-extraction routines. To more easily process diagonal-PASEF data we next built an automatic method detection algorithm into Spectronaut version 19 (**Figure 1, E**). Using this algorithm, Spectronaut will automatically determine any PASEF-based data-independent-acquisition type and apply the correct downstream processing, thus making seamless analysis of diagonal-PASEF acquisitions possible without any manual intervention of the user.

**Figure 1.**
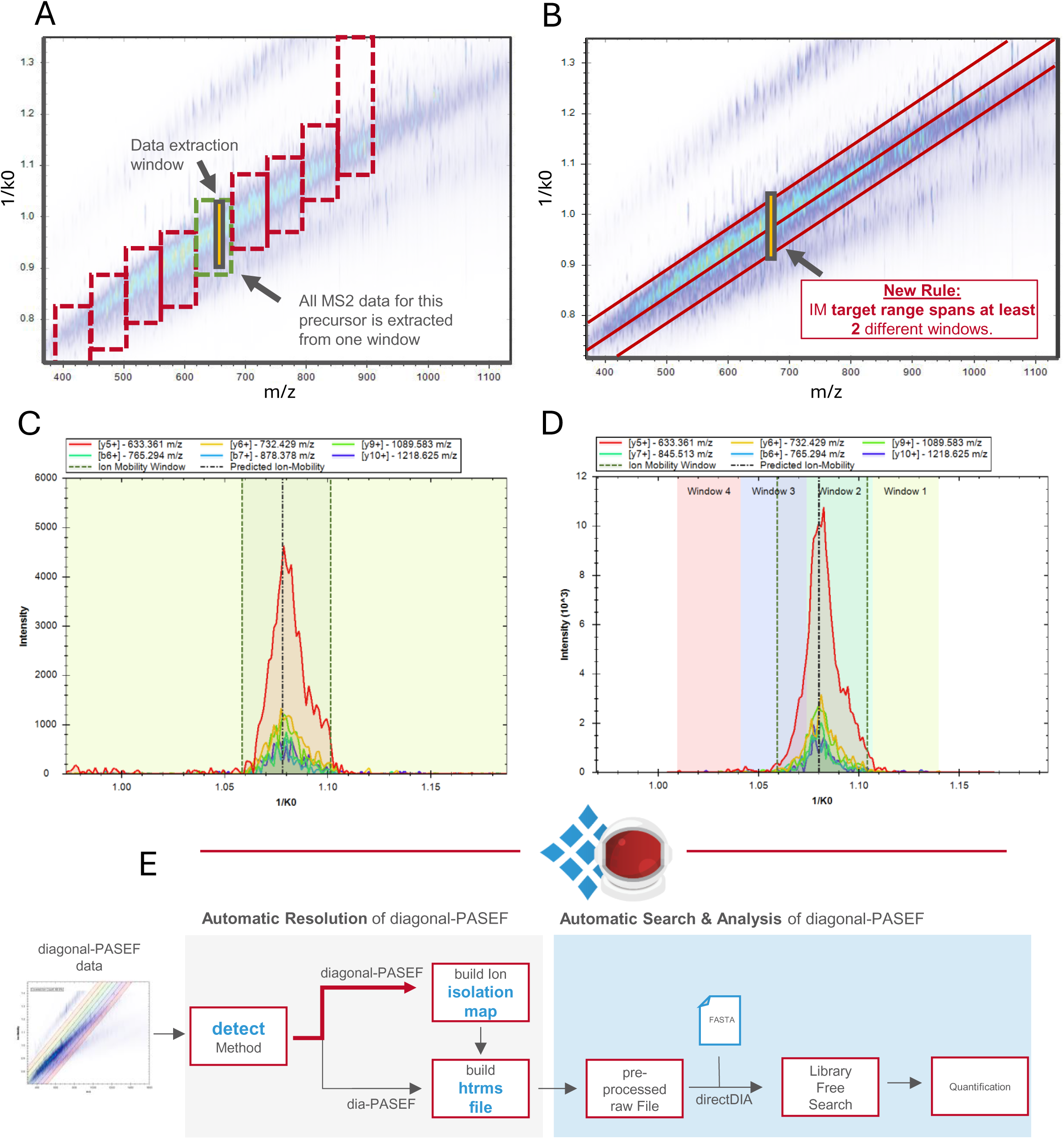
Spectronaut allows for the seamless integration of diagonal-PASEF workflows into existing analysis pipelines. A-B: Schematic overview of dia-PASEF (**A**) or diagonal-PASEF (**B**) acquisitions methods across an exemplarye ion distribution. Yellow areas indicate a schematic data extraction window designated for targeted MS2 data extraction. **C-D**: Extracted ion mobilogram (XIM) of a representative precursor “_DLYEDELVPLFEK_.2” on MS2-level from a dia-PASEF (**C**) or diagonal-PASEF (**D**) acquisition. Individual fragments are depicted as differentially colored lines. Data-extraction range is depicted as dashed vertical lines. Isolation windows governed by the acquisition method are shown as colored background. **E**: Simplified schematic visualization of a directDIA search and analysis workflow in Spectronaut 19.

To further optimize the processing of diagonal-PASEF data, we next investigated the suitability of the applied pulsar search & DIA-analysis settings which are applied during the directDIA pipeline (**Supplementary Figure 1**). For this we investigated the effect of the IM down-sampling by summation (IMsum) and DIA pre-processing settings on the precursor identifications for a representative diagonal-PASEF acquisition. We found that an IM summation value of 3 together with the DIA pre-processing set to “Legacy” mode improved the precursor, peptide and protein group identifications compared to the “BGS Factory Settings” which we deemed optimal for dia-PASEF acquisitions. During the “Legacy” pre-processing routine, SN19 will consider all micro-scans on the MS2 level that fall within the parent MS1-feature over the full IM width whereas in the “SN19 pre-processing” routine, only micro-scans that correspond to the IM-apex are considered. The usage of the MS2 apex micro scan only would significantly lower the number of ions used for the construction of the pseudo-DDA spectra during the spectrum centric search for diagonal-PASEF which is why we deviated from using this strategy.

Taken together, with the improved data-extraction of the Spectronaut algorithm we were able to upgrade the software to fully support various diagonal-PASEF acquisitions including synchro-and midia-PASEF while optimizing identification performance.

### Systematic optimization of diagonal-PASEF acquisitions for short gradients

Next, we investigated the performance of diagonal-PASEF in proteomics LCMS acquisitions in conjunction with our optimized Spectronaut analysis strategy. One of the key metrics of any MS-acquisition on the timsTOF systems is the cycle time of an acquisition method. The cycle time is determined by the number of PASEF ramps which are conducted at a defined ion mobility elution (or ramping) time. Here we used the timsTOF HT mass spectrometer and operated the system in a way that we kept the ramp and accumulation times at 100 ms. Given the selected chromatography which generated peaks with a full width at half the maximum of about 6-12 seconds depending on the gradient length we decided to generate diagonal-PASEF MS methods ranging from a single diagonal-PASEF slice to twenty diagonal-PASEF slices (**Figure 2, A**). To ensure comparability of the methods and allow for an optimal sampling of the ion-cloud we kept the width of the methods constant at 200 m/z for all tested methods. Due to this fixed overall method width, the resulting diagonal-PASEF method varied in the width of the individual slices and ranged from 200 m/z for the single-slice method to 10 m/z for the 20-slice method (**Figure 2, B**).

**Figure 2.**
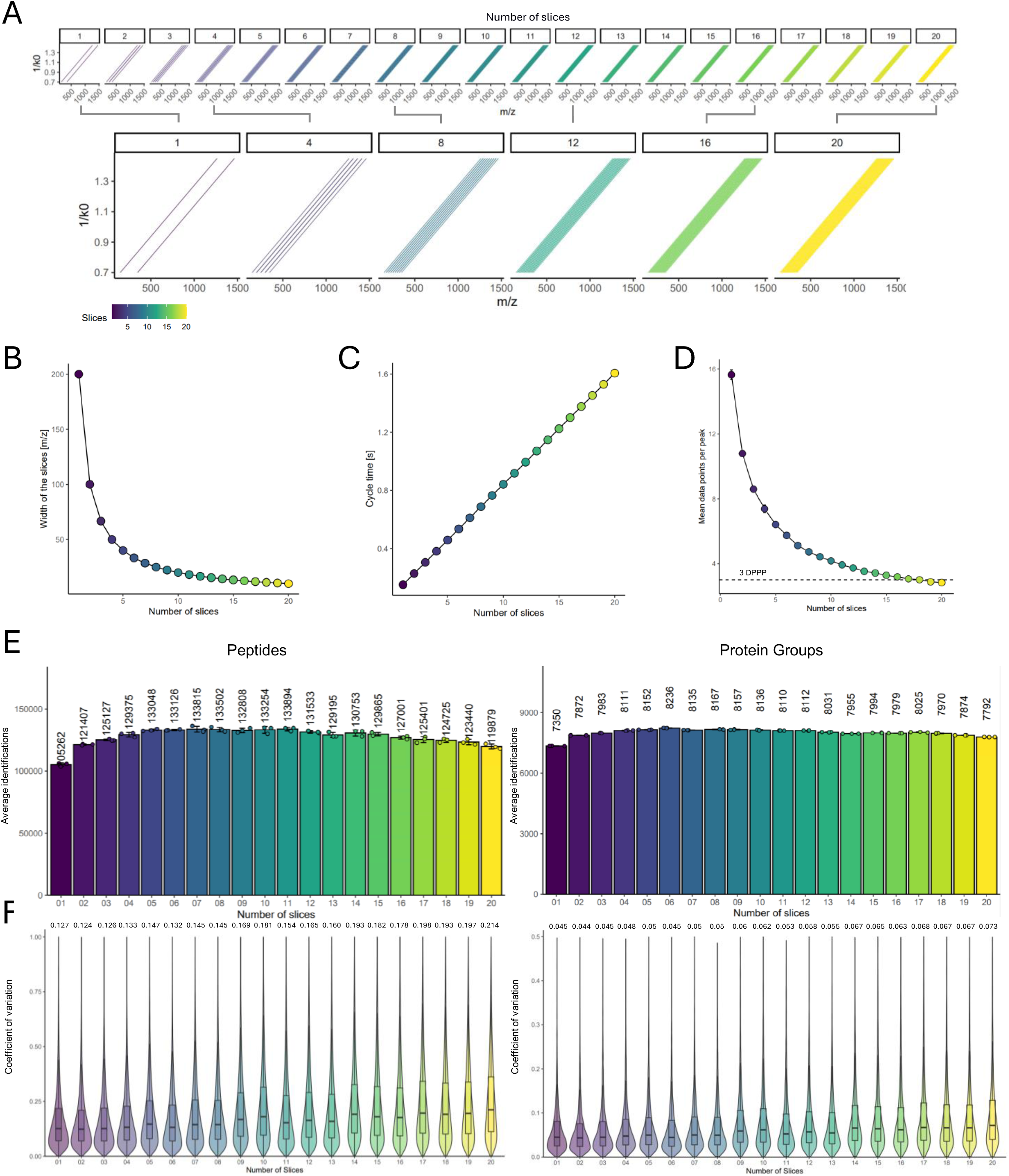
diagonal-PASEF measurements and data-analysis are stable across a wide range of tested methods. A: Depiction of tested diagonal-PASEF methods for method screening approach. Top: Methods ordered by ascending number of slices ranging from 1 to 20. All tested methods had an overall width of 200 m/z. Bottom: Magnification of selected methods. **B-C:** Width of slices in m/z (**B**) and resulting cycle time in s (**C**) for all tested methods from A. Each dot represents a single method. **D:** Data points per chromatographic peak (DPPP) of all tested methods from A in 17-minute gradients. Mean and standard deviation of mean shown (n = 3). Horizontal dashed line represents 3 DPPP. **E:** HeLa peptide (left) or protein group (right) identifications of all diagonal-PASEF methods from A in 17-minute gradients when 800 ng are injected. Mean identifications are shown. Error bars indicate standard deviation of mean (n = 3). **F:** Coefficient of variation (CV) of all peptide (left) or protein group (right) identifications from E. Boxplots indicate the inter-quartile range (IQR) from the lower quartile to the upper quartile. Central line indicates the median value of the population and whiskers indicate the 1.5 x IQR. Median CV is indicated above the plot. Outliers are not shown to aid the visual interpretation of the data.

The performance of diagonal-PASEF was examined by applying the generated MS methods in analytical triplicate injections of 800 ng of a HeLa sample in 17-minute gradients (∼37 samples per day) (**Figure 2 C-F**). The fastest cycling method consisting of a single slice had a cycle time of 0.15 seconds (**Figure 2 C**) and allowed to sample the chromatographic peak on average 15.6 times in 17-minute gradients (**Figure 2 D**). In comparison, the 20-slice diagonal-PASEF method had a cycle time of 1.61 seconds and achieved 2.8 data points per peak (DPPP) on average. We next investigated the peptide and protein group identifications that we obtained from the diagonal-PASEF acquisitions in directDIA searches in Spectronaut version 19 (**Figure 2 E-F**). Spectronaut processed all tested diagonal-PASEF acquisition seamlessly indicating that the software is suitable to analyze diagonal-PASEF methods with vastly different number of slices and sampling rates. We observed that the obtained average number of peptides varied strongly across the tested methods and gradient lengths (**Figure 2 E**). The method yielding the highest peptide identifications (133,815) consisted of 7 slices whereas most protein group identifications (8,236) were achieved when using a 6-slice method. For the peptide level the difference between the best-(7-slice) and worst-performing (1-slice) methods was 27.1% whereas on the protein group level this difference was only at 12.4%. These data indicated that diagonal-PASEF methods with different number of slices are more variable on the peptide level and to a lesser extent on protein level due to the coverage of proteins by multiple peptides.

Next, we investigated the quantitative precision of the diagonal-PASEF methods. For this we evaluated the median coefficient of variation (CV) of all methods that were acquired (**Figure 2 F, Supplementary Figure 1 A-B**). We observed that the median CV on the peptide level generally increased with an increase in the number of diagonal-PASEF slices from 12.7% at 1-slice to 21.4% with methods consisting of 20 diagonal-PASEF slices. On the protein group level, we observed similar trends and found the median CV to increase from 4.5% when using the 1-slice method to 7.3% when using the 20-slice method. When investigating the median CV over the DPPP we noticed that methods with >4 DPPP generally had a lower median CV compared to methods with <4 DPPP and that the median CV markedly increased with methods having more than 4 DPPP (**Supplementary Figure 2, C**). Given that the maximum peptide identifications with diagonal-PASEF methods were achieved when using 6-8 slices methods and that the lowest medium CV was achieved with methods having the least slices, we next investigated the identifications below 10% or 20% CV to identify the methods with the best quantitative precision (**Supplementary Figure 2, D**). This analysis revealed that a 6-slice diagonal-PASEF method achieved the highest quantitative precision with 91188 (68.5%) and 50837 (38.2%) of all peptides quantified with <20% and <10% CV, respectively. An increase in the number of DPPP did not yield more peptide identifications below 20% or 10% CV indicating that for diagonal-PASEF methods a balance between the number of diagonal-PASEF slices and the cycle time need to be reached to achieve best performance.

### Retention-time summation improves the signal-to-noise ratio of diagonal-PASEF acquisition and could boost the data quality

The data obtained so far in this study indicated that diagonal-PASEF methods generally result in very high quantitative precision which we attribute to the fast-sampling rate of the chromatographic peak of the methods. We therefore next sought to exploit this over-sampling and subjected a 2-slice diagonal-PASEF method to a retention time down-sampling strategy (**Figure 3 A**, **Supplementary Figure 3**). Similar strategies have previously been conceptualized for dia-PASEF but never been realized for diagonal-PASEF-type methods^12^. We developed the concept of retention-time down-sampling by summation (RTsum) subsequent to the ion mobility summation (IMsum) which is already conducted within the Spectronaut analysis pipeline. In this approach multiple dia-PASEF scans are down-sampled in the 1/k0 dimension to improve the signal-to-noise ratio of the derived XIM. In Spectronaut, RTsum can be applied during the pre-processing of the search & analysis pipeline of any dia-PASEF or diagonal-PASEF data. Commonly the RTsum value can be set as any strictly positive integer and determines the number of adjacent MS1 and MS2 scan events that are summarized during the processing of the run. During these summarizations the effective RT of the scans will be averaged, as such if a RTsum value of 2 is chosen for an acquisition that yields 8 DPPP then the resulting acquisition will be rendered to 4 DPPP whereby every 2^nd^ adjacent MS1 or MS2 scan is summarized into a novel summarized scan (**Figure 3 A**). We next validated the effects of RTsum on a 2-slice diagonal-PASEF acquisitions of 1000 ng of a HeLa sample in 17-minute gradients (n=4) and tested summation values ranging from 1 (no-summation) to 4 (4x summation, **Figure 3 B-G**). As expected, we found the number of MS1 (**Figure 3 B**) or MS2 (**Figure 3 C**) scans post analysis in Spectronaut to decrease in direct proportion to the defined RTsum value. We also identified the mean DPPP for all precursors identified in a single acquisition to decrease in direct proportion with the utilized RTsum value although a buffering effect was observed for an RTsum value of 3 and 4 (**Figure 3 D**). We reasoned that this buffering might arise from populational effects as not all precursors will have exactly the same peak width throughout the analytical gradient (**Supplementary Figure 3 A**, ^24,25^). To more accurately investigate the effect of RTsum on the peak shape we assessed the XIC and DPPP of the peptide-precursors FISADVHGIWSR (3+) and VIGFSPEEVESVHR (3+) (**Figure 3 E**, **Supplementary Figure 3 B-C**). Without RTsum, the 2-slice diagonal-PASEF method sampled the chromatographic peak of FISADVHGIWSR (3+): 15.2 and VIGFSPEEVESVHR (3+): 42 times on average across all four replicates. When applying an RTsum of 2, 3 or 4 the DPPP for FISADVHGIWSR (3+) were reduced to 9.25 (39.2% reduction), 5.75 (62.2% reduction) and 5.75 (62.2% reduction) respectively. For VIGFSPEEVESVHR (3+) applying an RTsum of 2, 3 or 4 reduced the DPPP to 24.2 (42.4%), 16 (61.9%) and 12.2 (71%) respectively. There was a significant correlation (p = 0.000000138; R^2^ = 0.99) between the measured and the theoretically expected DPPP after RT summation indicating that RTsum does not introduce DPPP artifacts (**Figure 3 F**). To validate that RTsum does not influence the peak-shape or chromatogram extraction we next investigated the XIC of the selected precursors and could indeed not observe any alterations of the XIC shape on the MS1 level for the mono-isotopic, M+1 or M+2 envelope (**Supplementary Figure 3 C**). In summary, these data provide evidence that the RT summation strategy provides the expected results and does not yield any sampling or XIC biases on the resulting data.

**Figure 3.**
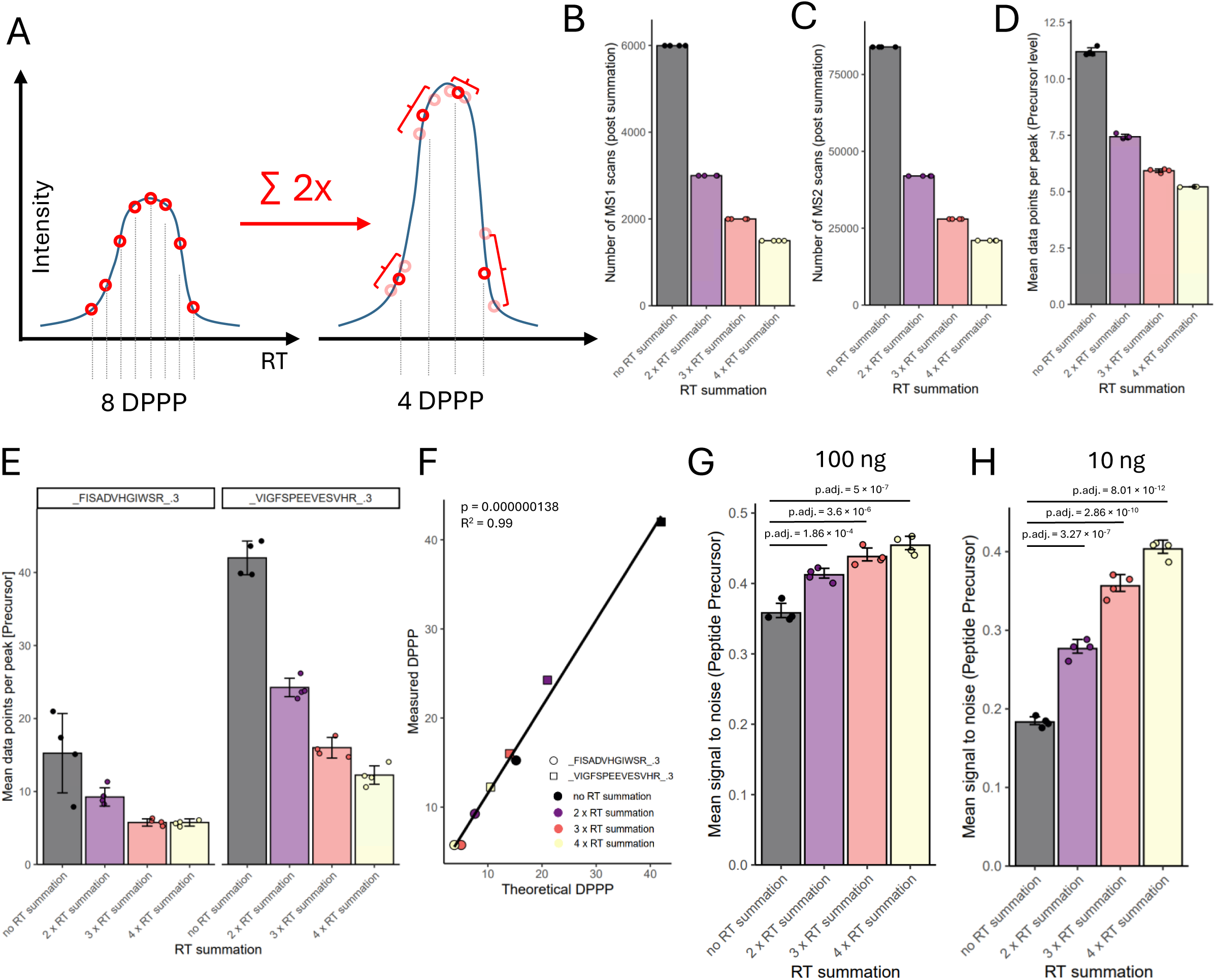
Retention time summation summarizes adjacent MS or MS/MS scans and thereby enhances the signal-to-noise ratio of diagonal-PASEF data. A: Schematic illustrating the concept of retention time summation. **B-D:** Characteristics of a 2-Slice diagonal-PASEF method subjected to 1-4 x retention time summation. Data from 1000 ng HEK-293 acquisitions in 17-minute gradients. **B:** Number of MS1 scans per injection post retention time summation. **C:** Number of MS2 scans per injection post retention time summation. **D:** Average median data points per chromatographic peak per injection post retention time summation shown. Error bars indicate standard deviation. **E:** Mean DPPP for selected precursors for a 2-slice diagonal-PASEF method with indicated retention time summation. Error bars indicate standard deviation of mean. Individual datapoints indicate replicate acquisitions. **C:** Spearman-based rank correlation of the empirically measured DPPP compared to the theoretically expected DPPP for the two selected precursors from E across tested retention time summation values. Diagonal line indicates the linear regression of the measured and theoretically determined DPPP. **G-H:** Mean signal-to-noise ratio on precursor level per injection shown post retention time summation for 100 ng (G) or 10 ng (H) input during the acquisition. Error bars indicate standard error of mean. Adjusted p-values from two-sided Tukey HSD test upon computation of one-sided ANOVA p-value (100ng: p = 5.27 x 10^-7^; 10ng: p = 1.69 x 10^-11^).

To investigate putative benefits of RTsum on the diagonal-PASEF data we investigated the average signal-to-noise ratio for all peptide precursors identified with a 2-slice diagonal-PASEF method (**Figure 3 G**). Strikingly we identified that using an RTsum of 2 (adj. p-val = 1.864 × 10^-4^), 3 (adj. p-val = 3.6 × 10^-6^) or 4 (adj. p-val = 5 × 10^-7^) provided a significantly higher signal-to-noise ratio compared to when not applying any RTsum (ANOVA p-value = 5.27 x 10^-7^) which we quantified by diving MS2 quantity of each precursor by its respective noise level (materials & methods). We confirmed these findings through injecting 10 ng of a HeLa material using the 2-slice diagonal-PASEF method and subjected the resulting data to RTsum of 2, 3 or 4 (**Figure 3 H**). We observed that RTsum significantly improved the signal-to-noise ratio for all detected peptide precursors compared to not applying any retention time down sampling by summation (ANOVA p-value = 1.69 x 10^-11^). In summary, these data indicate that RT summation might enhance the quality of the obtained data for diagonal-PASEF acquisitions especially for low-loadings.

### Retention-time summation improves the analytical performance of diagonal-PASEF acquisitions especially for low-inputs

We next sought to evaluate the performance of diagonal-PASEF in combination with RT summation and compared its performance to classic dia-PASEF acquisitions. For this we injected a HeLa digest at different loading amounts on our timsTOF HT mass spectrometer ranging from 5 ng to 1000 ng of net input material per acquisition (**Figure 4 A**) using diagonal-PASEF methods consisting of 1, 2, 4, 8 or 12 slices in quadruplicates (**Supplementary Figure 4 A-B**). To initially determine the optimal RT summation for each diagonal-PASEF method and loading we first analyzed each of the diagonal-PASEF acquisitions with an RT summation set to 1, 2, 3 or 4 (**Supplementary Figure 4 C**). We next investigated the average protein group and peptide identifications for each of these combinations as well as the average DPPP that the acquisitions would yield post RT summation (**Supplementary Figure 4 D**). As expected, RTsum enhanced the protein group and peptide identifications strongly for diagonal-PASEF methods consisting of 1, 2 or 4 slices across all tested loadings. We identified the strongest increase in the protein group and peptide identifications especially at low loadings. For example, at 5 ng of loading an RTsum value of 4 increased the protein group identifications by 21.4% and the peptide identifications by 34.2% compared to if no RTsum was applied for a 1-slice diagonal-PASEF method. RTsum-driven identification enhancements were observed at all loadings, but their extent faded with enhanced loading. For example, at 10ng and 100ng of loading applying an RTsum of 4 increased the peptide identifications by 20.4% and 1.5% respectively compared to no RTsum for a 1-slice diagonal-PASEF method. Interestingly, we noticed that RT summation generally proved to be detrimental for peptide or protein group identifications if applied to a diagonal-PASEF composed of 8 or 12 slices. For example, applying an RTsum of 2 on a diagonal-PASEF method with 12-slices reduced the peptide identifications across all tested loadings (5ng:-3.6%, 10ng:-2.0%, 100ng:-6.1%, 1000ng:-7.4%). We reasoned that if a diagonal-PASEF method already sampled the chromatographic peak between 3 and 5 times, this makes any RT summation incompatible with the chromatographic peak width. Based on these observations we decided to use the following RT summation values for our tested diagonal-PASEF methods across all loadings: 1-slice: 4x RTsum; 2-slice: 3x RTsum; 4-slice: 2x RTsum; 8-slice: 1x RTsum; 12-slice: 1x RTsum.

**Figure 4.**
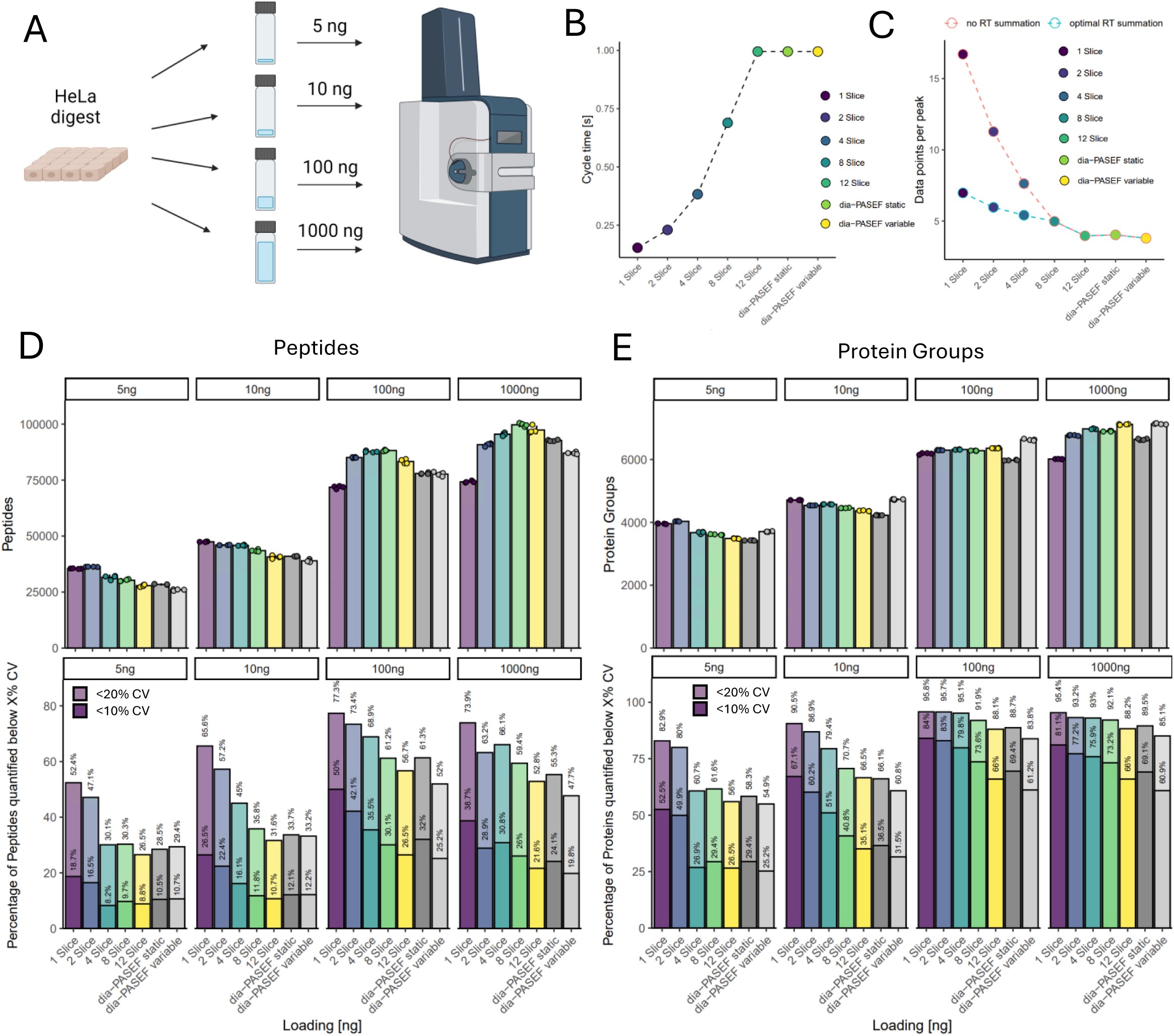
Retention time summation improves the overall identification and precision of diagonal-PASEF acquisitions. A: Schematic of experimental approach conducted to evaluate the performance of diagonal-PASEF against dia-PASEF across different tested loadings on the timsTOF HT. **B:** Cycle time of tested methods for loading ramp experiment depicted in A. **C:** Data points per peak of tested methods for loading ramp experiment pre (blue) or post (red) retention time summation. For the RT summation, the ideal value has been applied. **D-E:** Results of loading ramp from A. Top: Average peptide (D, left) or protein groups (E, right) identifications obtained for all tested methods across the indicated loadings. Error bars indicate standard deviation. Each datapoint indicates an individual acquisition. Bottom: Percentage of peptides (D, left) or protein groups (E, right) quantified below 10% or 20% coefficient of variation against overall identifications. Each bar represents the result of one search.

Next, we aimed at comparing the performance of diagonal-PASEF against classic dia-PASEF acquisitions. For this we acquired a dia-PASEF method composed of DIA windows with a static width (termed dia-PASEF static) which we computed using the timsControl method editor and a dia-PASEF method composed of dia-PASEF windows with variable widths (termed dia-PASEF variable) that we computed using an in-house software alongside all diagonal-PASEF acquisitions (**Figure 4 B-E, Supplementary Figure 4 B**). We designed both dia-PASEF methods to consist of exactly 12 PASEF ramps such that these methods would have the same cycle time and DPPP compared to the diagonal-PASEF method consisting of 12 slices (12 PASEF ramps, **Figure 4 B-C**). On average, the tested diagonal-PASEF methods yielded similar peptide and protein group identifications compared to the dia-PASEF control methods across all tested loadings (**Figure 4 D, E**). Diagonal-PASEF methods composed of <4 slices generally provided more peptide and protein group identifications at lower loadings. For example, at 5 ng of loading the 1-slice diagonal-PASEF method yielded 24.9% and 36.4% more peptides than the dia-PASEF static and variable methods respectively. On the protein group level, the same method yielded 15.6% and 6.7% more protein groups than the dia-PASEF static or variable methods respectively. In contrast, at the higher tested loadings of 100 and 1000 ng, the best diagonal-PASEF methods had more than 4 slices. The 12-slice diagonal-PASEF method yielded 2.8% or 18.3% more protein groups than the 1-slice method at 100 or 1000 ng respectively. This diagonal-PASEF method proved to be very competitive against dia-PASEF even at high loadings. At the highest tested loading of 1000 ng, the 12-slice method achieved 7.1% more protein groups and 5% more peptides than the dia-PASEF static method. Strikingly, the 12-slice diagonal-PASEF method only yielded 0.2% less protein groups but 11.8% more peptides than the highly optimized dia-PASEF variable method. We confirmed these results by evaluating the average number per peptides detected per protein group across all investigated methods (**Supplementary Figure 5**). In summary these data indicated that diagonal-PASEF are competitive against dia-PASEF methods across all tested loadings on the timsTOF HT.

Next, we investigated the quantitative precision of the tested dia-PASEF and diagonal-PASEF methodologies. For this we computed the protein groups or peptides quantified below 20% or 10% coefficient of variation (CV, **Supplementary Figure 6 A**) and analyzed the analytes quantified below a certain CV threshold relative to the overall identifications obtained (**Figure 4, D-E**). All tested diagonal-PASEF methods provided a very high quantitative precision. In general, we noticed that the relative quantitative precision was reduced with increasing number of diagonal-PASEF slices across the tested methods. For example, at 100 ng of loading 77.3% of all peptides were quantified below 20% CV using a 1-slice method whereas with the 12-slice method only 56.7% of all peptides were quantified below 20% CV (**Figure 4, D**). Despite this effect the tested diagonal-PASEF had a notably higher quantitative precision compared to the investigated dia-PASEF methods. At 5 ng of loading the average tested diagonal-PASEF method quantified 37.3% of all peptides and 68.2% of all proteins with less than 20% CV while the average dia-PASEF method quantified only 29.0% or 56.6% of all peptides or proteins respectively with less than 20% CV. The same pattern was observed for all other tested loadings. For example, on the peptide level the average diagonal-PASEF method quantified 47.0%, 67.6% or 63.1% of all peptides below 20% CV while the average dia-PASEF method quantified only 33.5% 56.7% or 51.5% of all peptides below 20% CV at a loading of 5, 10 or 100 ng. These trends did also hold up when quantifying the absolute number of analytes below 10% or 20% CV whereby diagonal-PASEF methods generally outcompeted dia-PASEF methodologies regardless of the utilized loading (**Supplementary Figure 6 A**). In summary, these data indicate that diagonal-PASEF methods not only provide higher overall identification rates than dia-PASEF methods but that these methods also allow for a better quantification of the detected analytes.

To dissect whether the higher number of quantified proteins or peptides did not arise due to overall higher identifications rates of diagonal-PASEF we next looked at the run-to-run correlation of all replicate injections for all methods and loadings (**Supplementary Figure 6 B**). As expected, we identified a higher average run-to-run correlation for the diagonal-PASEF compared to the dia-PASEF acquisitions at all tested loadings. All tested diagonal-PASEF method had an average run-to-run correlation on the peptide level as expressed by the Spearman-based rank correlation rho value of 0.902, 0.937, 0.972 and 0.963 at 5, 10, 100 and 1000 ng of loading respectively. In comparison, the dia-PASEF methods only achieved a correlation rho value of 0.887, 0.913, 0.961 and 0.948 at 5, 10, 100 and 1000 ng of loading respectively (**Supplementary Figure 6 C**). We generally observed all methods to achieve a better run-to-run correlation at higher loadings with the best correlation being obtained at 100 ng of loading (**Supplementary Figure 6 D**). These results were also confirmed on the protein group level although the detected differences were smaller, consistent with the observed trends for proteins and peptide CV in Figure 2F (**Supplementary Figure 6 E-G**). The largest difference in the run-to-run correlation was observed at 5 ng of loading. Here the 1-slice diagonal-PASEF method had the best run-to-run correlation with a spearman rho of 0.983 while that of dia-PASEF static was only 0.946. Interestingly when looking into individual run-to-run correlations we noticed that diagonal-PASEF appears to be more suited to precisely quantify especially low-abundant proteins (**Supplementary Figure 7 A, B**). Taken together these data indicate that diagonal-PASEF outperforms dia-PASEF in terms of quantitative precision and run-to-run correlation while frequently also yielding higher overall identifications

### diagonal-PASEF achieves a higher number of targets in controlled quantitative experiments compared to dia-PASEF

Given that the tested diagonal-PASEF methods in conjunction with RTsum yielded more peptide and protein identifications as well as higher quantitative precision compared to dia-PASEF we next evaluated the methods in more realistic scenarios. We and others have previously shown that quantitative controlled experiments (CQEs) are a suitable tool in assessing key metrics of analytical LCMS methods through the derivation of a ground truth^7^. Therefore, we set out and prepared a CQE in which we combined four different species (*H. sapiens*, *C. elegans*, *S. cerevisiae*, *E. Coli*) in different ratios (materials & methods) and prepared two distinct samples termed “A” and “B” which would yield predefined fold changes when measured together (**Figure 5 A**). We then acquired these samples at 5, 10, 100 and 1000 ng of loading on the timsTOF HT with a 17-minute analytical gradient using the dia-PASEF static and variable methods. We compared these dia-PASEF methods against a 12-slice diagonal-PASEF method which provided the overall highest protein group identifications at high loadings (≥ 100 ng) and a 2-slice diagonal-PASEF method which had the overall highest protein group identifications at low loadings (< 100ng). The resulting data were acquired in quadruplicates, and we investigated several key-parameters which we have previously shown to contribute to the analytical performance of LCMS methods in CQEs^7^. These include i) an identification metric for which we summarized the peptides or protein groups from samples A & B, ii) a precision metric for which we computed the coefficient of variation and iii) an accuracy metric for which we computed the median fold change error (**Figure 5 B, C**). First, we investigated the data on the peptide level (**Figure 5 B**). The 2-slice diagonal-PASEF method had the overall highest peptide identifications at a tested loading of 5 and 10 ng while at 100 ng and 1000 ng of loading the 12-slice diagonal-PASEF method achieved higher peptide identifications. At the highest tested loading of 1000 ng, the 12-slice diagonal-PASEF method achieved 2% more peptide identifications than the dia-PASEF variable method. These data confirm previous findings and indicate that diagonal-PASEF methods with a larger number of slices are generally more suited for larger loadings than methods with smaller number of slices (**Figure 4**). Next, we evaluated the quantitative precision of the tested LCMS methods. The 2-slice diagonal-PASEF method had the overall highest relative quantitative precision across all tested loadings (**Figure 5 B, Supplementary Figure 8 A**). At 5 ng of loading the 2-slice diagonal-PASEF method quantified 71.0% and 36.1% of all analytes with a CV below 20% and 10% respectively, whereas the dia-PASEF variable method only quantified 46.3% and 21.2% of all analytes with a CV below 20% and 10% respectively. The 12-slice diagonal-PASEF method had the overall worst quantitative precision at the lowest tested loading. At the highest tested loading of 1000 ng, the 2-slice method quantified 91.8% and 75.6% of all analytes below 20% and 10% CV whereas the dia-PASEF static method only quantified 83.1% and 57.6% of all analytes below 20% or 10% CV respectively. At the highest tested loading the 12-slice diagonal-PASEF method quantified 79.7% and 50.1% of all analytes below 20% and 10% respectively thereby leading to the most overall peptides quantified below 20% and 10% CV. These data indicate that the diagonal-PASEF method provide high peptide identifications at a higher quantitative precision compared to the tested dia-PASEF methods at all investigated loadings. Finally, we evaluated the absolute fold-change error that we computed as percentage of the deviation between the empirically determined fold change by the analytical method and the expected fold change based on the experimental design (see Material & Methods, **Figure 5 B, Supplementary Figure 8 B**). Across all tested loadings we identified the 12-slice diagonal-PASEF method to achieve the best quantitative accuracy with an average median fold-change error of 18.8% compared to 24.7%, 22.0% and 20.6% for the 2-slice, dia-PASEF static and dia-PASEF variable methods respectively. We noticed that the 2-slice diagonal-PASEF method appeared to have an overall worse quantitative accuracy compared to the 12-slice method. Despite these differences, all four methods achieved a comparable fold-change error indicating that all methods have a similar quantitative accuracy. Taken together these data indicate that on the peptide level diagonal-PASEF methods achieve higher identifications and a quantitative precision and a comparable quantitative accuracy compared to dia-PASEF methods.

**Figure 5.**
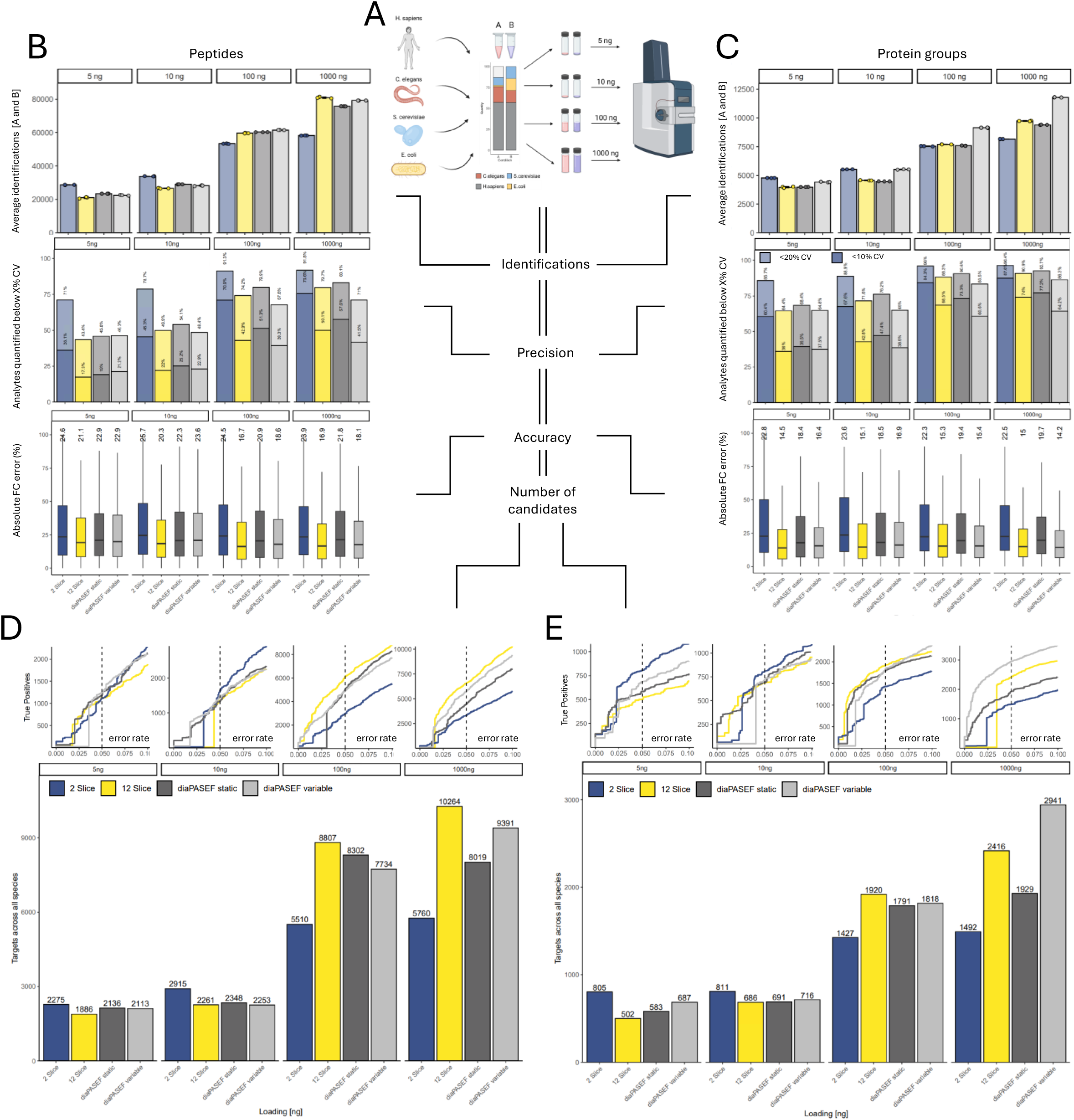
diagonal-PASEF and retention time summation outperform dia-PASEF acquisitions in quantitative controlled experiments. A: Schematic of the experimental approach conducted. **B,C:** Top: Average peptide (**B**) or protein group (**C**) identifications across samples A and B for all indicated diagonal-PASEF or dia-PASEF methods for the indicated loadings. Data points indicate individual replicates. Mean and standard deviation of mean shown. Middle: percentage of identifications below 20% CV (faint) or 10% CV (bold) as percentage of overall identifications. Percentage values above bars indicate the height of bars. Bottom: Absolute fold change error (%) peptide (**B**) or protein group (**C**) level for indicated methods and tested loadings. Median absolute fold change errors are indicated above the boxplots. Boxplots indicate the inter-quartile range (IQR) from the lower quartile to the upper quartile. Central line indicates the median value of the population and whiskers indicate the 1.5 x IQR. Median CV is indicated above the plot. Outliers are not shown to aid the visual interpretation of the data. **D-E:** Top: True positive (TP) identifications over the error rate of candidates shown for diagonal-PASEF and dia-PASEF methods on peptide (**D**) or protein group (**C**) level. Dashed vertical line indicates an error rate of 5% at which the number of true positives was read out. Bottom: Number of candidates with error rate < 5% for indicated method and loading.

An important aim of proteomics projects is to detect differentially abundant proteins or peptides which arise from, for example, statistical testing of biological conditions. We therefore next investigated the number of true candidates below 5% error rate for all tested loadings and acquisition methods to provide a more complete evaluation of the CQE (**Figure 5 D**). For this we performed pair-wise testing of all peptide identifications between samples A and B, assessed whether the analytes followed the projected fold-change trajectory and counted the number of correctly quantified true candidates below an error rate of 5% (see Material & Methods). The diagonal-PASEF methods achieved more candidates on the peptide level across all tested loadings compared to the dia-PASEF methods. At 5 ng and 10 ng of loading the 2-slice method allowed to identify 2275 and 2915 candidates respectively which was 6.5% or 21.1% more targets than the best performing dia-PASEF method (dia-PASEF static) achieved at these loadings. At 100 and 1000 ng of loading the 12-slice diagonal-PASEF identified the most candidates with 8807 and 10264 true positive identifications respectively which was 6.0% and 9.0% more than the best performing dia-PASEF method achieved. We confirmed these findings by computing the partial area under the curve (pAUC) for the true positives versus candidate list plots below an error rate of 5% (**Supplementary Figure 9 A**).

Next, we repeated these analyses for all tested diagonal-PASEF and dia-PASEF methods on the protein group level (**Figure 5 C, E; Supplementary Figure 8, B, C, Supplementary Figure 9 B**). On the identification level we found the dia-PASEF variable method to achieve the overall highest identifications at 100 and 1000 ng of loading in the CQE. Notably, at a loading of 5 and 10 ng, the 2-slice diagonal-PASEF method achieved more protein group identifications. When investigating the quantitative precision, we again identified the 2-slice diagonal-PASEF method to quantify most analytes below 20% or 10% across all tested loadings, similar to the data previously obtained for the peptide level (**Figure 5 B**). In agreement with this, the median fold-change error was also observed to be similar for all tested methods. As on the peptide-level data the 12-slice diagonal-PASEF error achieved the lowest average median fold change error across all loadings of 15% whereas the dia-PASEF variable, static and the 2-slice diagonal-PASEF method had a median fold change error of 15.7%, 19% and 22.8% respectively. We next computed the true candidates retained below an error rate of 5% (**Figure 5 E**) on the protein group level. Similar to the peptide-level data we found the 2-slice diagonal-PASEF method to achieve the most targets at low loadings with 805 and 811 candidate identifications for a loading of 5 and 10 ng respectively. At higher loadings the 12-slice diagonal-PASEF method outcompeted the 2-slice method and achieved 1920 and 2416 candidate identifications at 100 and 1000 ng of loading respectively. While the 12-slice diagonal-PASEF method identified 5.6% more candidates at 100 ng compared to the dia-PASEF variable method it identified 17.9% less candidates at 1000 ng of loading. In summary, these data indicate that at the protein level diagonal-PASEF methodologies outcompetes dia-PASEF methods up to 100 ng of loading but that at higher loadings, dia-PASEF still achieves superior candidate recovery.

## Discussion

In this study, we present a novel data analysis strategy for fast-cycling diagonal-PASEF, utilizing retention time-based down-sampling by summation of scans, implemented in Spectronaut 19. We comprehensively optimized several diagonal-PASEF method parameters. In the human cell line HeLa, diagonal-PASEF resulted in higher peptide identifications and similar protein identifications compared to dia-PASEF. Importantly, the precision at both the peptide and protein levels was consistently higher with diagonal-PASEF than with dia-PASEF. In a quantitative controlled experiment, diagonal-PASEF yielded similar accuracy in fold-change measurements but generally identified a higher number of true positive peptides at an error rate of 5% (based on the t-test results), compared to traditional dia-PASEF.

We found that diagonal-PASEF can efficiently cover the main ion distribution of the TIMS elution with as few as one to three slices. Consequently, these methods can have very short cycle times (∼0.05 – 0.5 seconds), allowing for oversampling of chromatographic peaks that typically elute over several seconds (∼3 – 9s). To optimally analyze oversampled data, we summed corresponding consecutive MS1 and MS2 scans over the RT dimension. Methods with fewer slices, i.e., shorter cycle times, allowed for a higher degree of down-sampling compared to methods with more slices. Methods with the highest number of slices (8, 12) were incompatible with down-sampling, as the resulting method cycle time would not sufficiently sample the chromatographic peaks. When optimizing diagonal-PASEF, we observed that for lower sample amounts (5 and 10 ng), methods with fewer slices were superior to those with higher slice numbers (Figure 4D and E). This could be due to the observed increase in signal-to-noise ratio for methods where retention time down-sampling is applied (Figure 3G). Improvements in the signal-to-noise ratio likely lead to better identification and higher precision for low sample amounts, compared to methods without retention time down-sampling. We reason that in such cases, many signals might be close to the noise floor, making the signal-to-noise ratio a limiting factor for the search engine to translate spectra into peptide identifications. For HeLa, we observed that diagonal-PASEF generally achieved higher peptide identifications than dia-PASEF. At the protein level, identifications were similar for both methods, leading to better sequence coverage of the identified proteins and, consequently, more information on peptide-specific traits such as PTMs, isoforms, and proteoforms. Strikingly, we found the quantitative precision of peptides and proteins to be consistently higher for diagonal-PASEF compared to dia-PASEF, confirming recent data from another study^18^. These findings render diagonal-PASEF particularly well-suited for studies aiming at peptide-level data, such as post-translational modifications (PTMs), peptidomics-based approaches, and immunopeptidomics.

To evaluate the performance of diagonal-PASEF for detecting differentially abundant analytes, we performed a controlled quantitative experiment and benchmarked it against dia-PASEF. We found peptide identifications to be similar across all tested sample loading amounts between the best diagonal-PASEF and dia-PASEF methods. In contrast to the observations from the single human proteome experiments using HeLa, at the highest sample loading, variable dia-PASEF yielded the most protein identifications. CQEs rely on mixing multiple proteomes and are thus significantly more complex.

Currently, this higher complexity seems to be handled more efficiently by the dia-PASEF data analysis pipeline. However, CQEs do not reflect the reality for most proteomics projects, which commonly comprise only one organism of interest. The variable dia-PASEF method and the diagonal-PASEF method with similar isolation widths had similar accuracy. Interestingly, we found the 2-slice diagonal-PASEF method to achieve the lowest accuracy, possibly due to the high number of interferences caused by the wide quadrupole isolation window. We observed no dependency of accuracy on peptide loading amount, indicating that diagonal-PASEF does not generally improve accuracy over dia-PASEF. When performing statistical testing for differential abundance, accuracy, precision and identifications are relevant. We found diagonal-PASEF to yield more true positive peptides and proteins that are statistically differentially abundant for most sample loading amounts compared to dia-PASEF. The increase in statistical power is explained by the increase in precision while maintaining accuracy. As with the HeLa peptide loading ramp, for low peptide loading amounts, diagonal-PASEF with few slices performed better, and at higher peptide loads, the method with many slices proved superior. Finally, variable dia-PASEF candidate recovery performance was best at the loading of 1000 ng, correlating with the highest identification of variable dia-PASEF.

Future improvements for diagonal-PASEF could include acquisitions with very narrow slices. In this study, we kept the overall width of the diagonal-PASEF method constant at 200 m/z. Other studies have shown that modulating this parameter could enhance the performance of diagonal-PASEF^18^. Therefore, in a more systematic assessment, the width of the diagonal-PASEF method could be adjusted against parameters such as the ramping time of the TIMS cells or the number of diagonal-PASEF slices to identify the optimal configuration. Additionally, the instrument control software for ion mobility and quadrupole scanning dynamics is being actively developed to enable new acquisition modes in the future, including variable slice methods that have recently been proposed ^18^. On the data analysis side, additional algorithms like the proposed precursor slicing^22^ and fingerprint midia-PASEF type of analyses^23^ could increase both identification and quantification performance.

Despite being a nascent technology, diagonal-PASEF already outperforms dia-PASEF in key metrics such as peptide identification and quantification. This is of particular interest for workflows focused on post-translational modifications, peptidomics, and immunopeptidomics, where higher quality peptide-level data can dramatically improve results. We expect this novel acquisition method to have the potential to replace dia-PASEF in the near future.

## Material & Methods

### Protein material from cell pellets

HeLa cell pellets were purchased from Cell Line Services. *E. coli* and *S. cerevisiae* digests were kindly provided by Dr. Audrey van Drogen*. C. elegans* Bristol strain worm digests were kindly provided by Prof. Monica Gotta (University of Geneva). Frozen HeLa cell pellets were purchased from Dundee cell products.

### Sample Preparation

HeLa cell pellets (Cell Line Services) were diluted in 100 µL of Biognosys lysis buffer and subsequently lysed and homogenized using a LE220Rsc ultrasonicator (Covaris). Following sonication, samples were denatured at 95°C for 5 minutes. All subsequent liquid handling and sample processing were automated using a Hamilton Microlab STAR system, equipped with a HMotion robotic arm, Hamilton heater shakers, and integrated third-party devices, including an Epoch microplate reader (Agilent), Ultraseal™ Pro plate sealer (Porvair Sciences), Xpeel® plate peeler (Azenta), and Ultravap® Mistral blow-down evaporator (Porvair Sciences). Protein concentrations were determined using the BCA assay (Thermo Fisher Scientific) according to the manufacturer’s instructions. For further processing, 75 µg of protein was used in a protein aggregation capture (PAC) protocol, as described previously ^3,27^. Peptide desalting for mass spectrometry was performed using an Oasis HLB μElution Plate (30 μm, Waters) following the manufacturer’s protocol. Desalted peptides were dried using a blow-down evaporator and reconstituted in LC solvent A (1% acetonitrile in water with 0.1% formic acid) containing Biognosys’ iRT-peptide mix for retention time calibration. Peptide concentrations in mass spectrometry-ready samples were measured using the mBCA assay (Thermo Fisher Scientific). For loading ramp experiments, the peptide digest was diluted in LC Solvent A to a final concentration permitting the infusion of 1 µL sample material per acquisition.

Mixed proteome samples for quantitative controlled experiment were prepared as previously described^7^. In brief, the *C. elegans* worms were first lysed using a bead mill (Eppendorf) upon suspension in 8 M urea and 0.1 M ammonium bicarbonates and thereafter subjected to preparation using the filter aided sample preparation protocol following by desalting using the Oasis HLB μElution Plate (30 μm, Waters). *E. Coli,* HeLa and *S. cerevisiae* samples were re-suspended in 8 M urea lysis buffer, vortexed and the protein concentration was determined using the BCA assay (ThermoFisher) according to the manufacturer’s instructions. Samples were then reduced (10 mM Tris(2-carboxyethyl)phosphine) and alkylated (40 mM CAA Chloroacetamide) at 800 rpm, 37°C for 60 minutes followed by a digestion with trypsin (Promega) in a 1:50 ratio overnight at 800 rpm, 37°C. Prior to digestion the 8 M urea buffer was diluted using 0.1 M ammonium bicarbonate to dilute the urea to 2 M. Upon digestion, samples were desalted using the HLB 96 well-plates (Waters) according to the manufacturer’s instructions and peptide concentration was determined using the mBCA kit (ThermoFisher Scientific). Proteomes were mixed in the following ratios*: H. Sapiens:E.coli:S.cervisiae:C.elegans* at 1:10:2.1:0.77. For loading ramp experiments, the peptide digest was diluted in LC Solvent A to a final concentration permitting the infusion of 1 µL sample material per acquisition.

### Liquid chromatography coupled mass spectrometry

Unless otherwise indicated, 800 ng of peptides were injected to an IonOpticks Aurora series Ultimate CSI 75 µm x 250 mm C18 reversed phase column (AUR3-25075C18-CSI) on a Thermo Scientific™ EASY-nLC™1200 nano-liquid chromatography system connected to a Bruker Daltonics timsTOF HT mass spectrometer equipped with a Captive Spray II ion source. LC solvents were A: water with 0.1% FA; B: 80 % acetonitrile, 0.1% FA in water. The nonlinear LC gradient was 1 – 45% solvent B in 18 minutes followed by a column washing step in 90% B for 2.5 minutes, and a final equilibration step of 1% B for 3 minutes at 60°C with a flow rate set to a ramp between 600 to 400 nL/min (min 0: 600 nL/min, min 18: 400 nL/min, washing at 600 nL/min) as follows: Time[min]:%B[%]:Analytical Flow[µL/min]: 0:1.0:0.6, 1:6.0:0.60, 2:8.0:0.6, 3:10.0:0.6, 4.16:11:0.58, 5.33:13:0.57, 6.5:15:0.56, 7.7:16.0:0.53, 8.83:18:0.52, 10:20.0:0.511, 11.33:21:0.49, 18:45:0.4.For HeLa or mixed proteome loading ramp experiments the indicated loading amount was injected onto the analytical separation column. All acquisitions were conducted in positive ionization mode at a source capillary voltage of 1500 V. During the operation of the instrument, every acquisition had in-batch calibration enabled whereby the Tuning MIX ES-TOF CCS reference list was used as calibration template. Automatic calibration was conducted on calibrants with m/z of 622.029, 922.0098, 1221.9006 (Chip Cube High Mass Reference kit, Agilent Technologies).

### Method design of dia-PASEF and diagonal-PASEF methods

All dia-PASEF and diagonal-PASEF methods consisted of one full range MS1 scan from 100 –1700 m/z with an applied ion mobility range from 0.7 –1.45 1/k_0_ with ramp and accumulation times set to 70 ms (100 % duty cycle). Default processing and TIMS settings were applied for all methods, the mobility detection threshold was set to 5000 and no denoising was applied to any acquisition. For all acquisitions the applied collision energy was set to a custom ramp between 0.8 and 1.45 1/k_0_ as follows: [1/k_0_:eV]: 0.8:30, 0.95:30, 1.05:35, 1.15:45, 1.25:55, 1.35:65, 1.45:55. The ‘Base’ type was used for any stepping in the custom gradient.

The’dia-PASEF static’ method was generated using the timsControl dia-PASEF Window Editor (timsControl V6.06, Bruker Daltonics GmbH & Co. KG) and consisted of 12 dia-PASEF ramps resolved over 36 boxes with a width set to 55 Da and a mass overlap set to 0.5 Th between adjacent boxes. The effective m/z coverage of dia-PASEF boxes was 190.5 – 1444.5 and the method resulted in a cycle time of 0.99 s.

The’dia-PASEF variable’ method was generated using an in-house software (Biognosys AG) and consisted of 12 dia-PASEF ramps resolved over 50 boxes with variable width and a mass overlap set to 0.5 Th between adjacent boxes. The effective m/z coverage of dia-PASEF boxes was 175 – 1450 and the method resulted in a cycle time of 0.99 s.

The diagonal-PASEF methods were generated using the timsControl diagonal-PASEF Window Editor (timsControl V6.06, Bruker Daltonics GmbH & Co. KG). For all HeLa and mixed proteome loading ramp experiments the width of the diagonal-PASEF beam was set to 200 m/z and resolved by 1, 2, 4, 8 or 12 slices leading to an effective isolation width per slice of 200, 100, 50, 25, 16.67 m/z. The resulting m/z start and end ranges were constant for all diagonal-PASEF methods and resulted in an m/z start of the slices between 375 and 1250 m/z and an m/z end of 567 m/z and 1447 m/z. All diagonal-PASEF methods had an effective overlap of the slices of 1 Th as previously described elsewhere ^23^. For the implementation of the overlap between slices, a midia-PASEF licence was enabled and utilized in the timsControl software. The cycle time of the methods varied depending on the number of diagonal-PASEF slices. For the ramping of the diagonal-PASEF slices, the diagonal-PASEF slices varied between 1 and 20 slices, and an overlap of the slices was disabled for this experiment.

### Diagonal-PASEF calibration

All diagonal-PASEF methods were calibrated as per the recommendations by Bruker. In brief, upon successful calibration of the m/z and the mobility of the timsTOF HT each diagonal-PASEF method was loaded, and the ESI-L Low Concentration Tuning Mix (Agilent Technologies) was infused at 3 µL/min through the ESI Apollo source. The diagonal-PASEF quadrupole calibration routine was conducted, and the methods were calibrated on the Background ions while the ions from the calibration mix were excluded through the Tuning Mix ES (ESI) exclusion list. Diagonal-PASEF methods were calibrated on a bi-weekly basis to ensure sufficient isolation of the quadrupole.

### Mass spectrometric data analysis

All dia-PASEF and diagonal-PASEF data were analyzed using Spectronaut version 19.8 (Biognosys AG, Schlieren, Switzerland). The directDIA pipeline was used for all acquisitions and conditions were subjected to individual searches whereby all replicates belonging to one search were grouped. For all analyses the modified default settings were used as described by the manufacturer (Biognosys AG). In brief, Carbamidomethylation (C) was used as fixed modification and Acetyl (Protein-N-term) and Oxidation (M) were used as variable modifications. The PSM, Peptide and Protein group false discovery rates (FDR) were set to 1% during the pulsar search. No imputation was applied during the quantification of the analytes and minor (peptide) grouping was conducted on Stripped Sequence level. The following modifications were applied: For dia-PASEF acquisitions the dia-PASEF pre-processing mode “Automatic” was used, IM sampling reduction was set to 7 and RT sampling reduction was set to 1. For diagonal-PASEF acquisitions, the dia-PASEF pre-processing mode “Legacy (Spectronaut 18)” was used, IM sampling reduction was set to 3 and RT sampling reduction was set to the indicated value in the main text (1-4). For the generation of representative XIM plots a 100 ng HeLa injection conducted with dia-PASEF or diagonal-PASEF was utilized and the data searched by setting the IM summation to 1 and dia-PASEF pre-processing set to “fast SN19” for dia-PASEF and diagonal-PASEF. For the systematic testing of the optimal search and analysis settings for diagonal-PASEF data the values were used as indicated in the main text. FASTA files were generated using the UniProt sequence database, the following FASTA files were used: For HeLa acquisitions: *Homo sapiens* Uniprot/Swiss-Prot database from 2024-07-01 containing 20435 entries, for mixed proteome acquisitions: *Homo sapiens* Uniprot/Swiss-Prot database from 2024-01-01 containing 20418 entries, *Saccharomyces cerevisiae* (strain ATCC 204508 / S288c) Uniprot/Swiss-Prot database from 2024-01-01 containing 6727 entries, *Escherichia coli* (strain K12) Uniprot/Swiss-Prot database from 2024-01-01 containing 4530 entries, *Caenorhabditis elegans* Uniprot/TrEMBL databased from 2024-01-01 containing 27453 entries. For all FASTA files in mixed proteome experiments, only the peptide sequences that are unique to each species were retained.

## Statistical Data Analysis

All data analyses were conducted in the R programming language using the tidyverse package environment for all data handling and modification^28,29^. Prior to data handling, the quantitative data was extracted on peptide precursor level from Spectronaut using a custom report schema. Correlation analyses were conducted curing the cor.test() function in R and “Spearman” or “Pearson”-type correlation were applied depending on the normality of the datasets as indicated^30^. Normality of the data was assessed through the Shapiro.test() function in R^31^. For the computation of any DPPP value we utilized the “EG.DatapointsPerPeak” column from Spectronaut. For the computation of the signal-to-noise ratio we divided the quantity (“EG.Quantity”, Spectronaut) value of each precursor by the noise (“EG.Noise”, Spectronaut) value. The number of MS1 or MS2 spectra were obtained from the columns “R.MS1Spectra” or “R.MS2Spectra” from Spectronaut. One-sided ANOVA was conducted using the aov() function in R and pairwise comparisons were conducted using Tukey’s ‘Honest Significance Difference’ method through the TukeyHSD() function in R. Quantitative peptide precursor level data was exported from Spectronaut including for the CQE experiments. Subsequently, using R the data was collapsed to the desired quantitative unit (peptide stripped sequence or protein groups). The data were normalized such that the median of the stable human background proteome was equal for all acquisitions. Next, quantitative units, that have zero variance were removed. The quantitative data was log-transformed. Statistical analysis was performed on normalized MS2 quantitative level using a two-sided t-test assuming equal variance. FDR was calculated using the Storey approach^32^. The fold change was calculated based on log2(condition 2) / log2 (condition 1). Next, the ground truth was annotated, human as “background” and the other organisms as “spike-in” with a theoretical fold change column. Subsequently, the data was sorted by ascending p-value and then the true positives and true negatives were counted. All the regulated candidate proteins are sorted by p-value and categorized as true and false positives based on the ground truth pipetting. Finally, number of true candidates can be calculated based on error rate of finding the control species.

## Author Contributions

C.B., R.B., and L.R. designed the project. S.K.S. and J.K. supported the experimental design of the research. R.B., L.RA, S.S. prepared the samples. C.B. and I.K. designed the acquisition methods and carried out the measurements. C.B., O.M.B., B.S. and R.B. performed the data analysis. O.M.B, S.W. and E.C. wrote the software. C.B., S.K.S. and R.B. wrote the paper. R.B., T.G. and L.R. supervised the project. All authors critically revised the manuscript and approved its content.

## Data availability

The raw MS data, the spectral libraries and the quantitative data tables have been deposited to the ProteomeXchange Consortium via the PRIDE partner repository with the dataset identifier PXDxxxxxxxx. The Saved projects from Spectronaut can be viewed with the Spectronaut Viewer (www.biognosys.com/spectronaut-viewer).

## Abbreviations

CQE: controlled quantitative experiment
DIA: data-independent acquisition
DDA: data-dependent acquisitions
DPPP: data points per peak
FDR: False discovery rate
IM: Ion mobility
LCMS: Liquid chromatography mass spectrometry
PASEF: parallel accumulation serial fragmentation
RT: retention time
RTsum: retention time down-sampling by summation
TIMS: trapped ion mobility spectrometry
TOF: time of flight
XIC: Extracted ion chromatogram
XIM: Extracted ion mobilogram

## Acknowledgements

We would like to thank Oliver Raether and Christopher Krisp for their input into the experimental design and assistance in the planning of the project. We would like to thank Daniel Hornburg and Nigel Beaton for his review of and input to the manuscript.

---

**Supplementary Figure 1:**
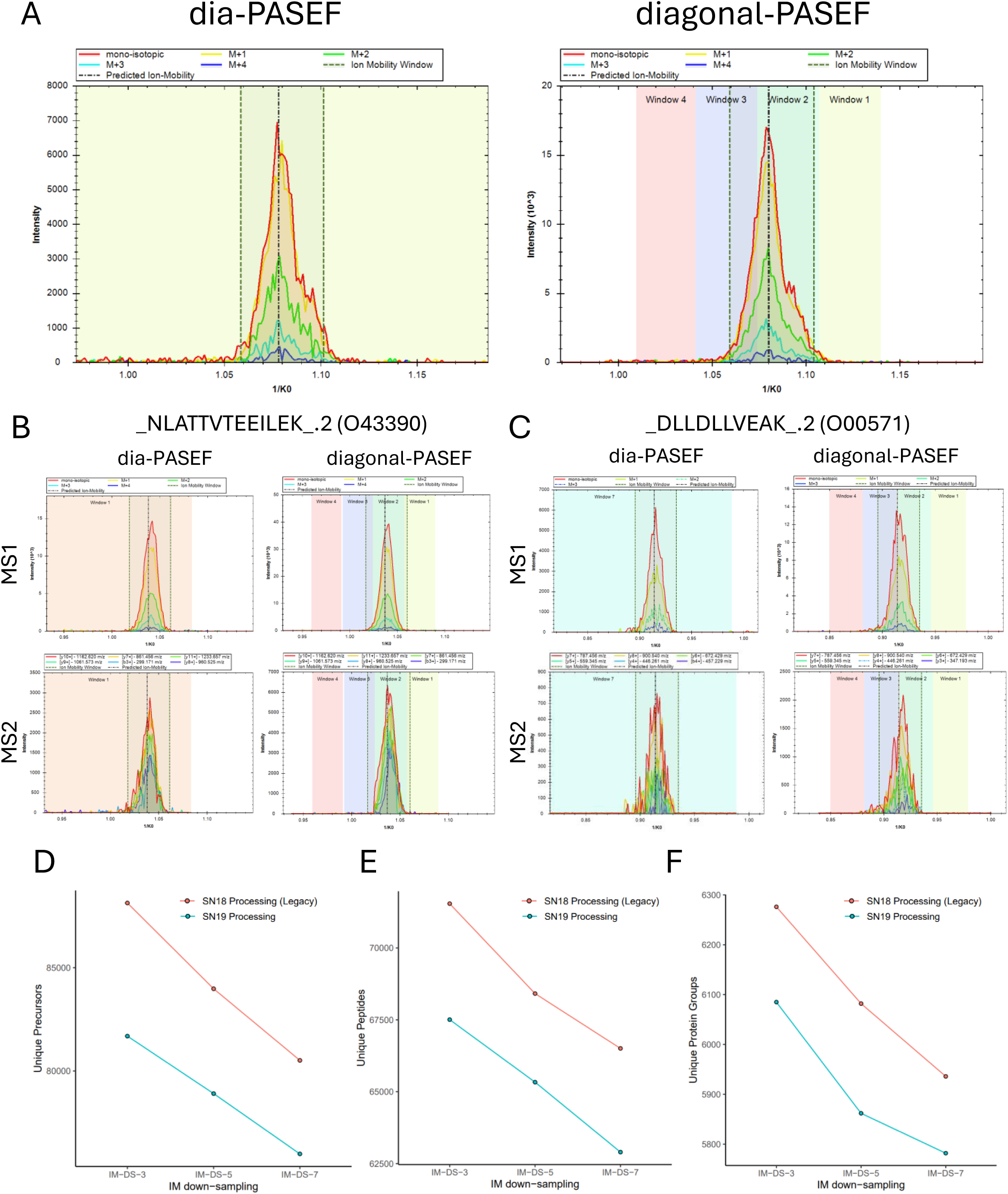
A: Extracted ion mobilogram (XIM) of a representative precursor “_DLYEDELVPLFEK_.2” on MS1-level from a dia-PASEF (left) or diagonal-PASEF (right) acquisition. Isotopic envelope depicted as differentially colored lines. Data-extraction range is depicted as dashed vertical line. Isolation windows governed by the acquisition method are shown as colored background. **B-C:** Extracted ion mobilogram for precursor “_NLATTVTEEILEK_.2 “ (**B**) and “_DLLDLLVEAK_.2” on MS1 (top) or MS2 (bottom) level from dia-PASEF (left) or diagonal-PASEF (right) acquisitions. Individual fragments are depicted as differentially colored lines. Data-extraction range is depicted as dashed vertical line. Isolation windows governed by the acquisition method are shown as colored background. **D-F:** Optimization of SN parameters “DIA pre-processing” and “Ion Mobility down-sampling” for a 100 ng HeLa acquisition using 2-slice diagonal-PASEF method at 17-minute analytical gradients on precursor (**D**), peptide (**E**) or protein group (**F**) levels.

**Supplementary Figure 2.**
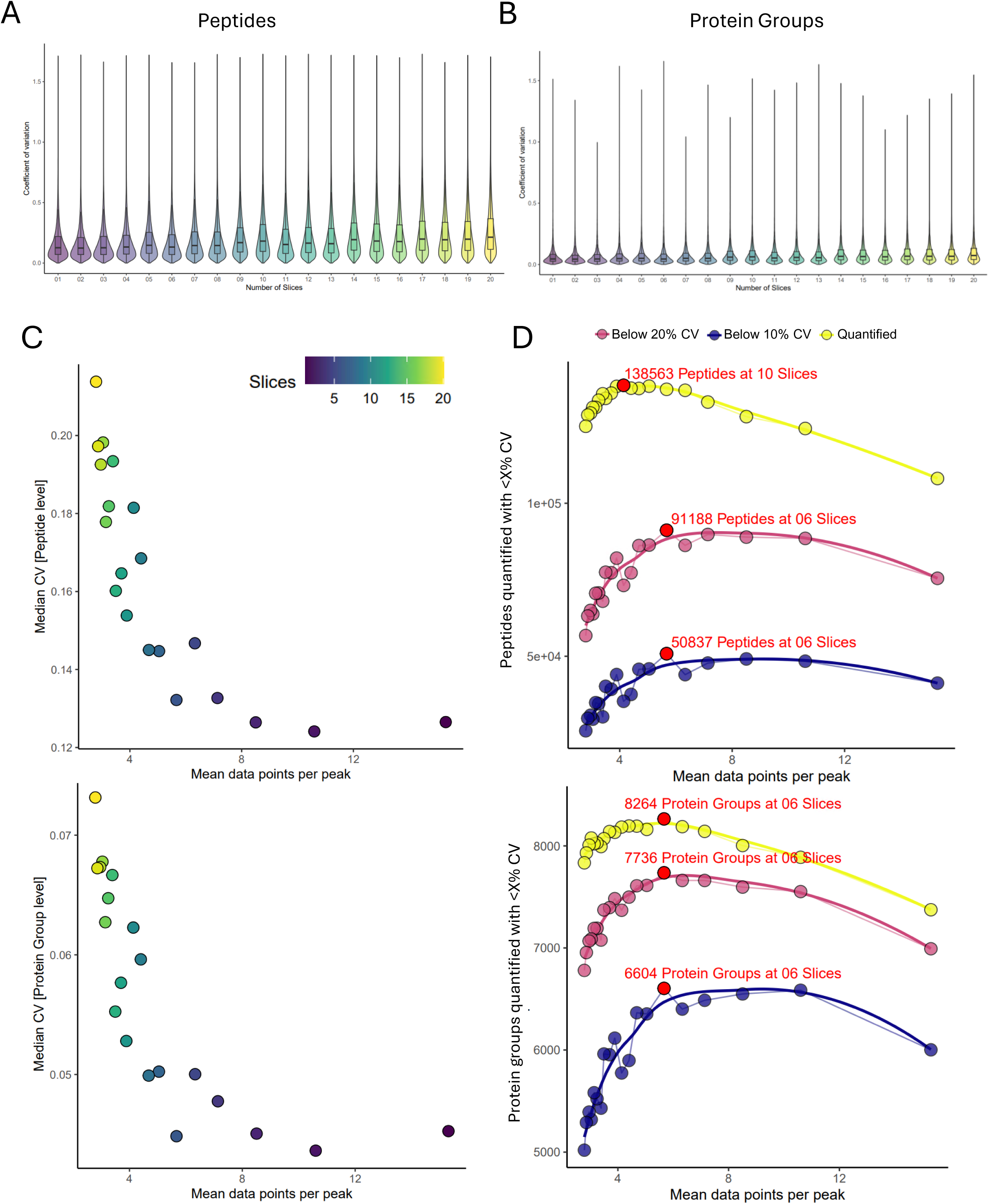
A-B: Coefficient of variation (CV) of all peptide (left, A) or protein group (right, B) identifications for all tested diagonal-PASEF methods. Boxplots indicate the inter-quartile range (IQR) from the lower quartile to the upper quartile. Central line indicates the median value of the population and whiskers indicate the 1.5 x IQR. Median CV is indicated above the plot. Outliers are not shown to aid the visual interpretation of the data. **C**: Scatter plot of mean data points per peak against the median coefficient of variation computed on the peptide (top) or protein group (bottom) level. Each data point depicts an individual diagonal-PASEF method. **D**: Mean data points per peak of each method against the peptides (top) or protein groups (bottom) quantified, quantified below 20% CV or 10% CV. The best performing method is indicated in red. Each data point represents an individual method.

**Supplementary Figure 3.**
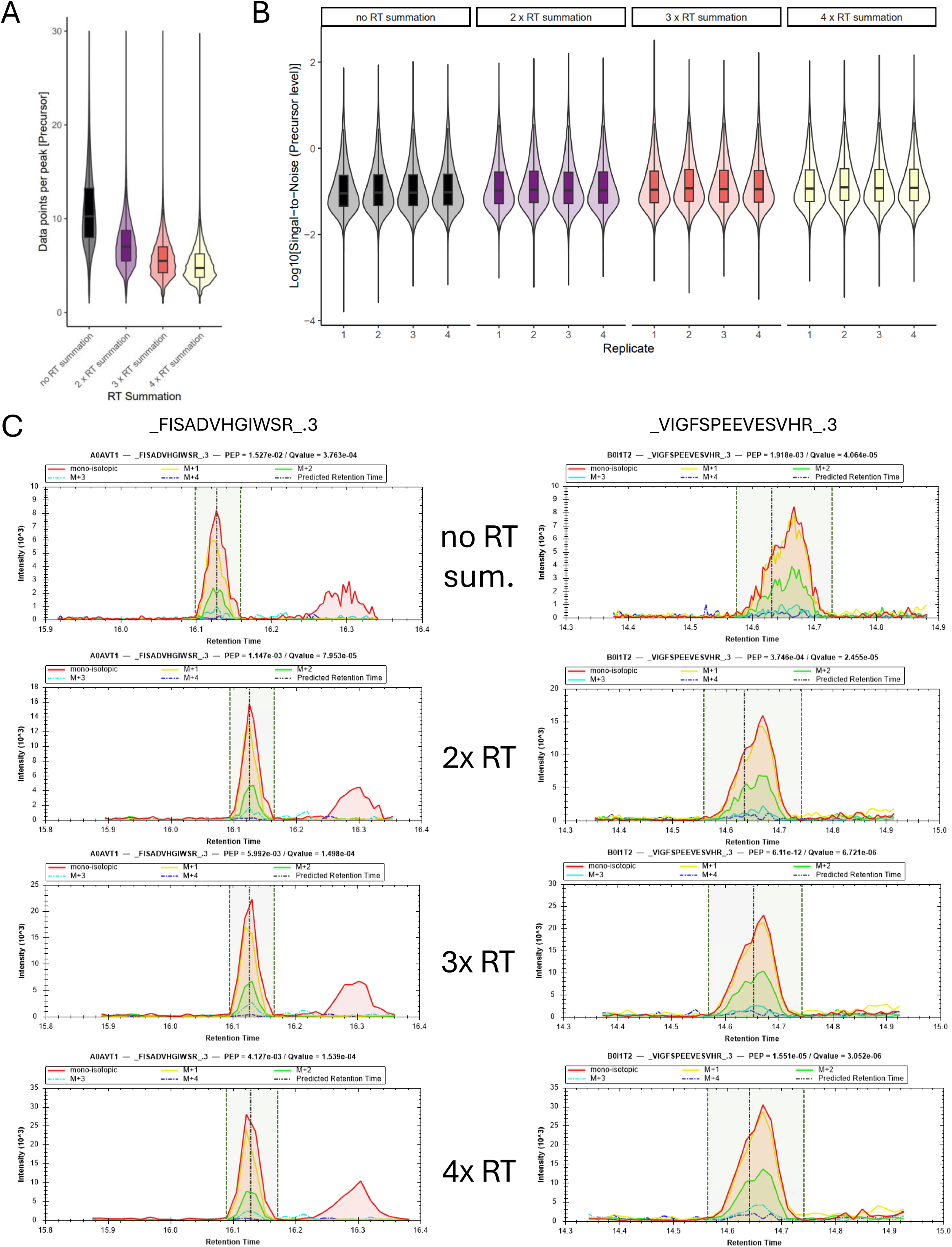
A: Data point per peak distribution on precursor level for a 2-slice diagonal-PASEF method post indicated retention time summation. Mean DPPP for each precursors across all replicate shown**. B:** Log10 signal-to-noise ratio for all detected precursors of the method from A across different retention time summation values. Individual boxplots and violin plots depict different replicates. **C:** Extracted ion chromatogram (XIC) of two selected precursors identified using the same method as from A-B subjected to the indicated retention time summation. All Boxplots indicate the inter-quartile range (IQR) from the lower quartile to the upper quartile. Central line indicates the median value of the population and whiskers indicate the 1.5 x IQR. Median CV is indicated above the plot. Outliers are not shown to aid the visual interpretation of the data.

**Supplementary Figure 4.**
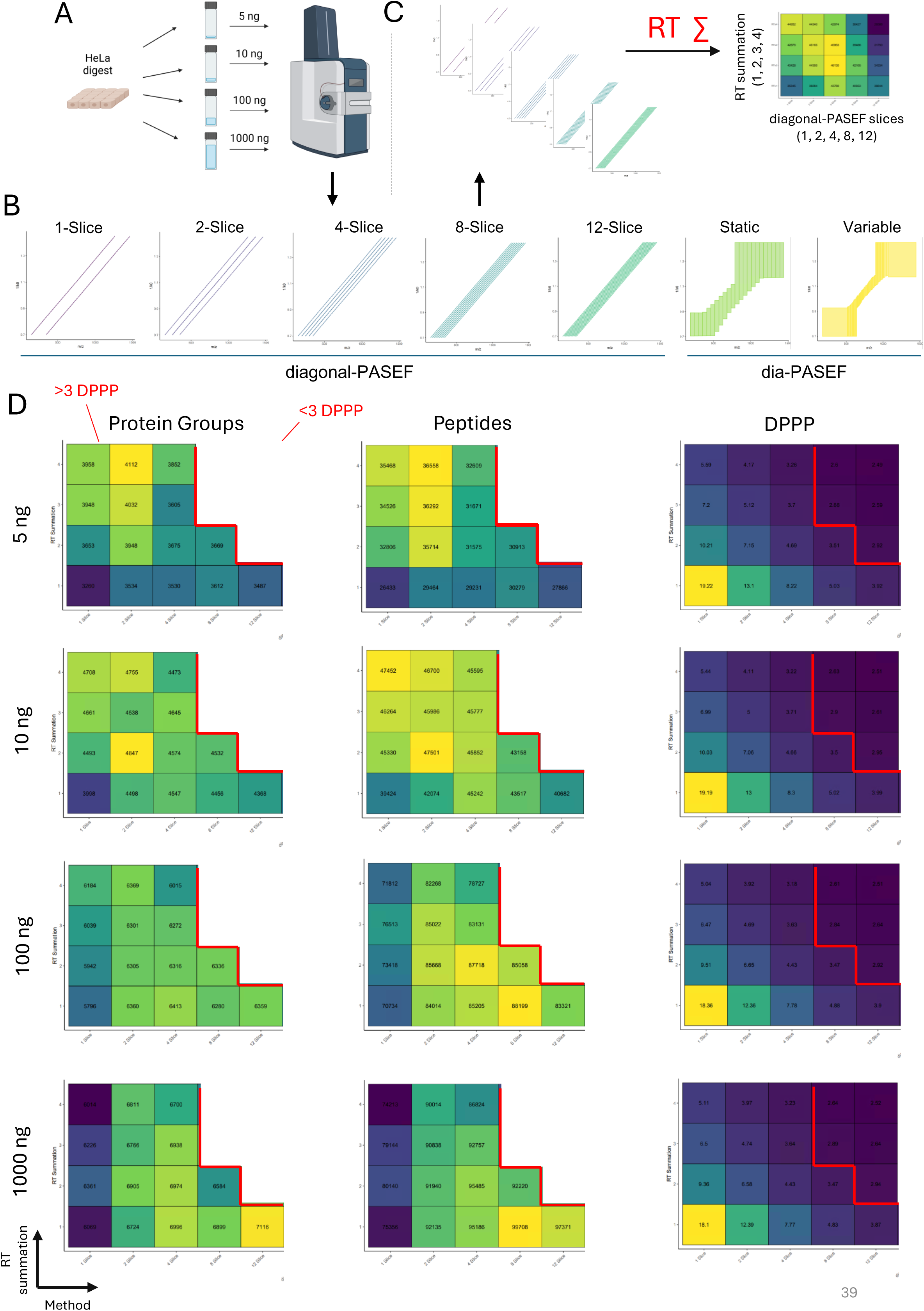
A-C: Summary of the experimental approach conducted to benchmark the retention time summation. **A**: Depiction of acquisition strategy, **B:** Overview over the individual diagonal-PASEF methods that were acquired; **C**: Overview over the systematic resting of retention time summation from 1-4 for all indicated methods. **D**: Results from systematic testing of the retention time summation. Average protein groups (left), precursors (center) and data points per peak (right) across four replicates are shown for each tested diagonal-PASEF method (x-axis) against the utilized retention time summation value (y-axis). Results are shown for the indicated loading from 5 (top) to 1000 ng (bottom). Red line indicates whether a tested method – RT summation combination achieved more or less than three DPPP. Black-frames indicate the RT summation values that achieved the highest precursor identifications for each utilized diagonal-PASEF method (center).

**Supplementary Figure 5.**
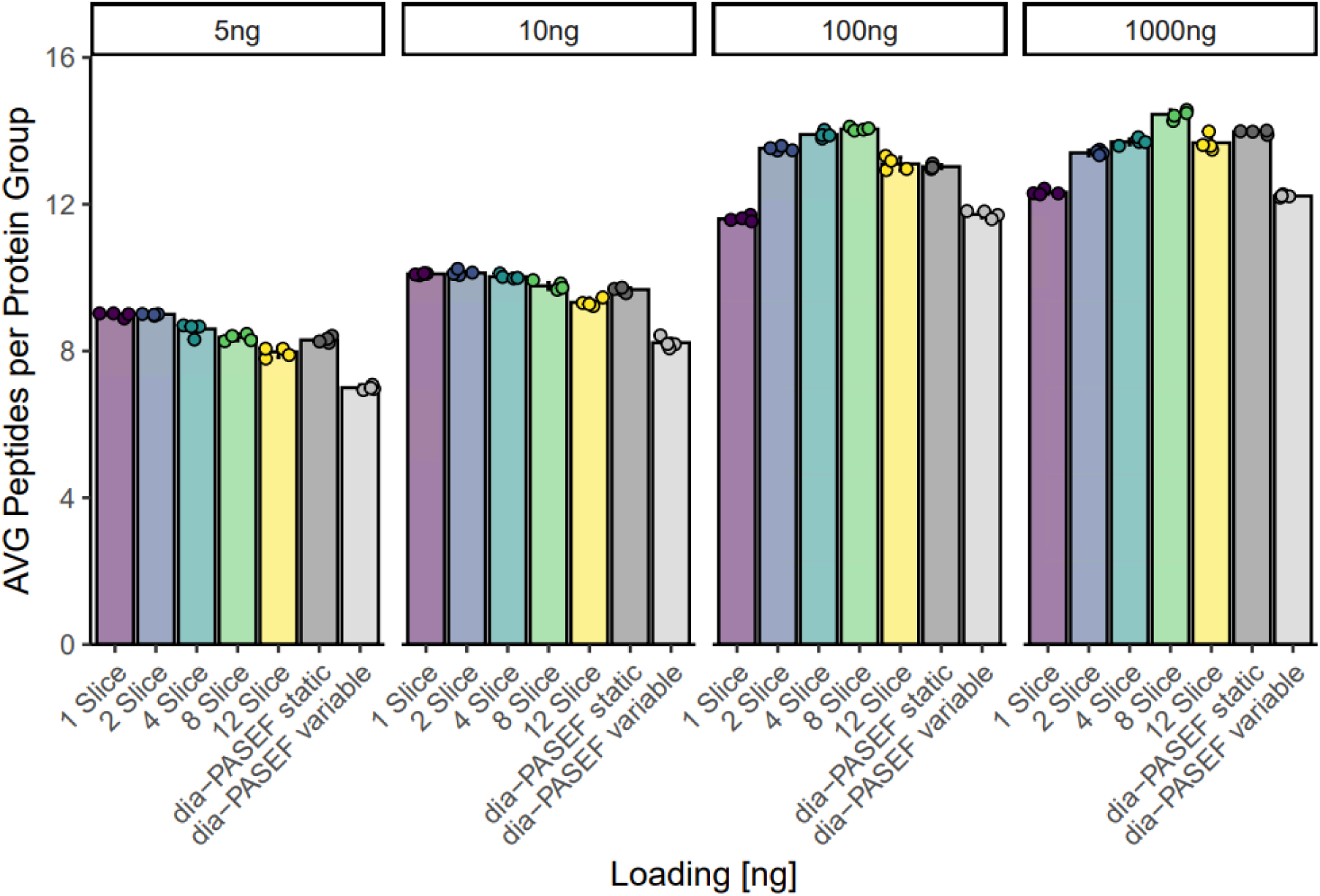
Average number of peptide identifications per protein group for all tested dia-PASEF and diagonal-PASEF methods for the indicated loadings. The height of the bar represents the average value across quadruplicates with standard deviation around mean shown. Data point indicate individual replicates.

**Supplementary Figure 6.**
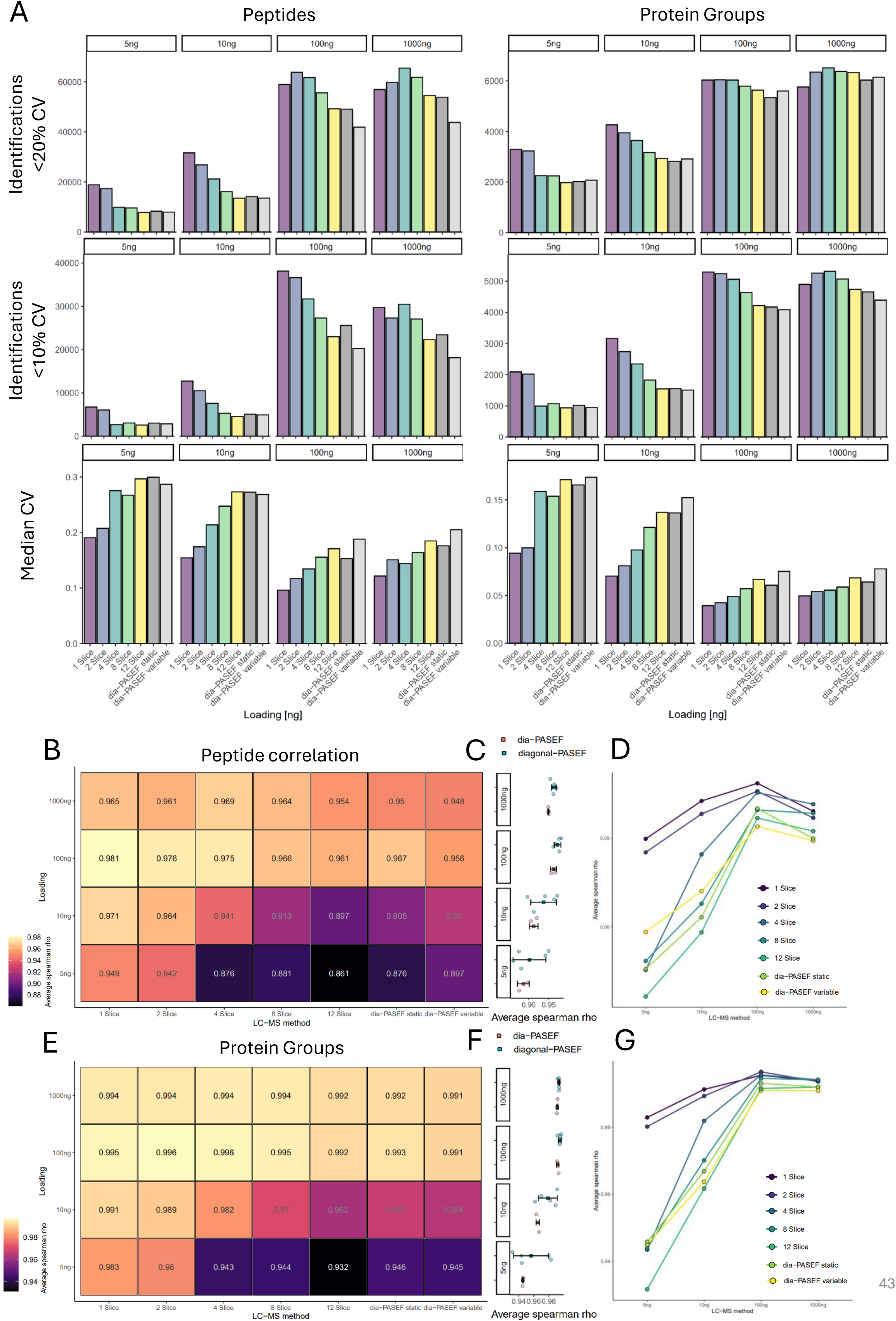
A: Results from the loading ramp experiment for all tested diagonal-or dia-PASEF methods. Protein group (left) or precursor (right) identifications below 20% CV (top) or 10% CV (center) or median CV (bottom) are shown. **B:** Average Pearson correlation on protein group level of all replicates against each other for each tested diagonal-PASEF method (x-axis) and loading (y-axis). **C:** Average person correlation from B stratified by acquisition-type. Central datapoint indicates the mean correlation score for each acquisition type. Error bars indicate standard deviation around mean. Individual datapoints indicate individual methods. **D:** Average person correlation from B visualized across different loadings. Individual acquisitions are color coded. E-G: Same analyses as depicted in B-D but on precursor level. For all panels, the optimal retention time summation was applied to all diagonal-PASEF acquisitions as follows: 1-Slice: 4xRTs, 2-Slice: 3xRTs, 4-Slice: 2xRTs, 8-Slice: 1xRTs, 12-Slice: 1xRTs.

**Supplementary Figure 7.**
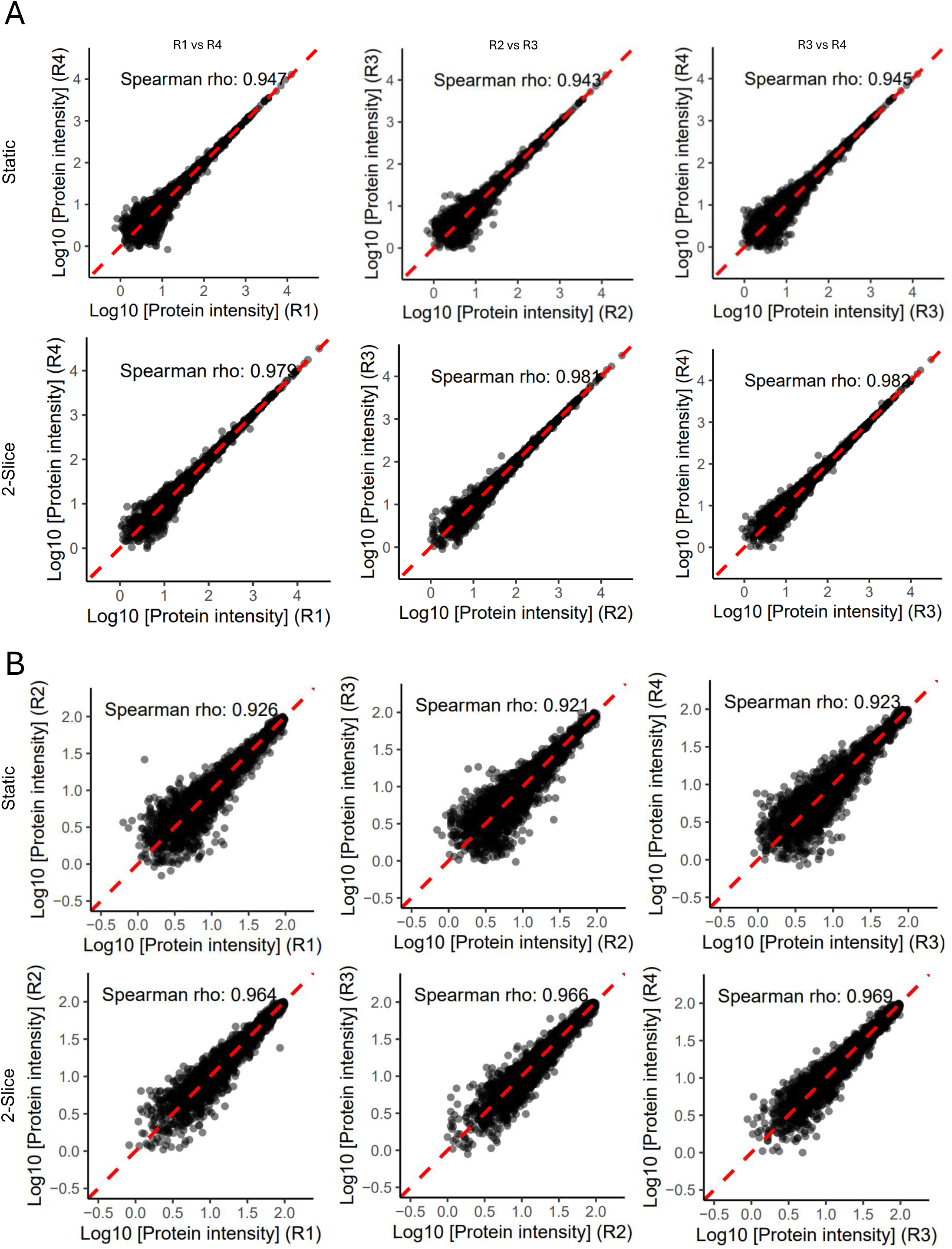
A: Representative correlations of individual replicates for the dia-PASEF static (top) or the 2-slice diagonal-PASEF method on the protein level for 5 ng HEK-293 acquisitions. The 2-slice diagonal-PASEF method was subjected to a retention time summation of 3. Each data-point represents a unique protein. Dashed red line represents the linear regression between the two indicated replicates. Correlation values from spearman-based rank correlation. **B:** Same as in A but for proteins with log10(protein quantity) <= 2.

**Supplementary Figure 8:**
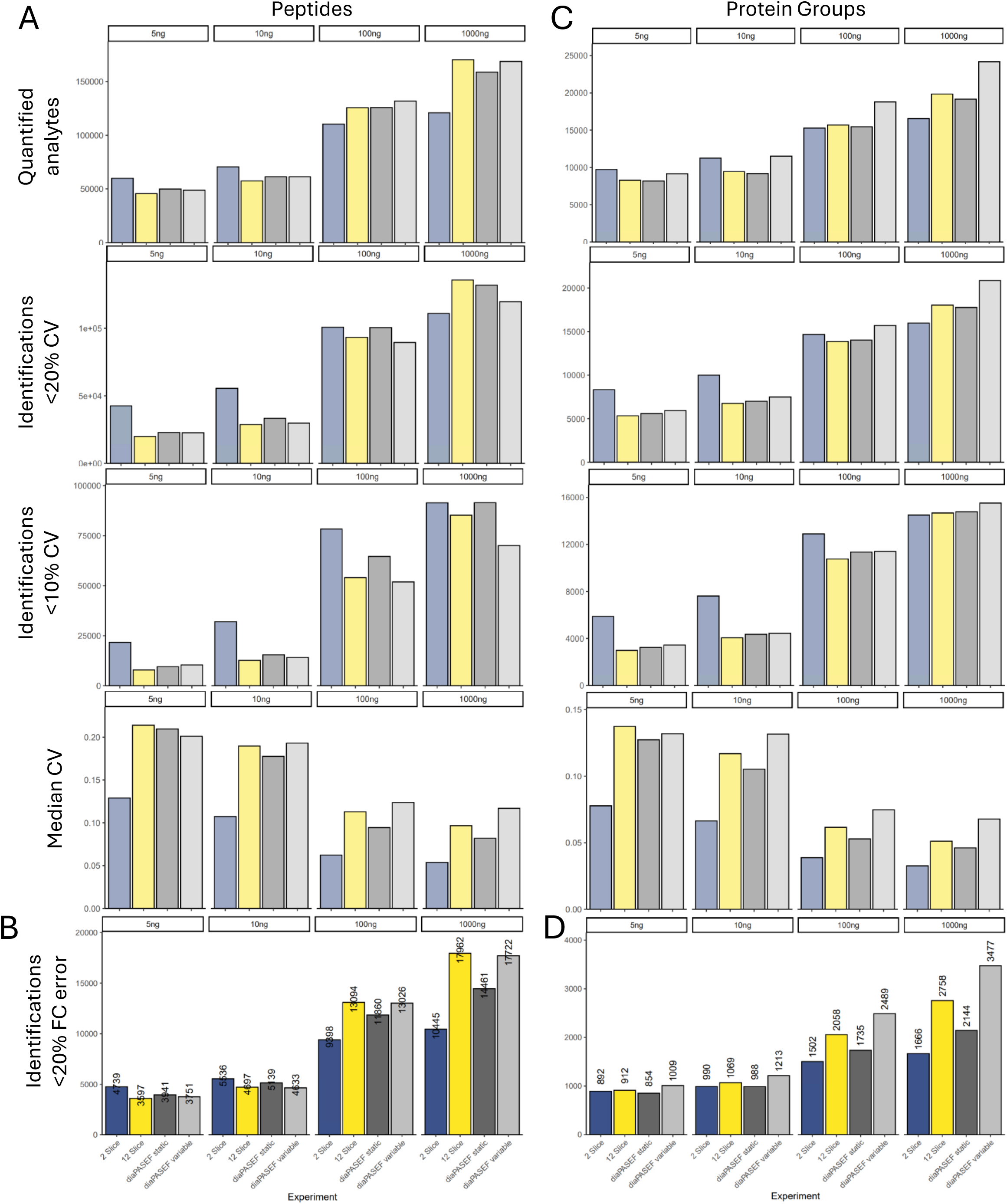
A, C: Number of overall quantified analytes (top), analytes quantified below 20% CV (middle) and below 10% CV (bottom) for all tested diagonal-PASEF and dia-PASEF methods for the indicated loadings on peptide-(**A**) or protein group (**C**) level. **B, D:** Number of peptides (**B**) or protein groups (**D**) quantified with a fold-change error of less than 20%. Values indicate the height of the bar.

**Supplementary Figure 9:**
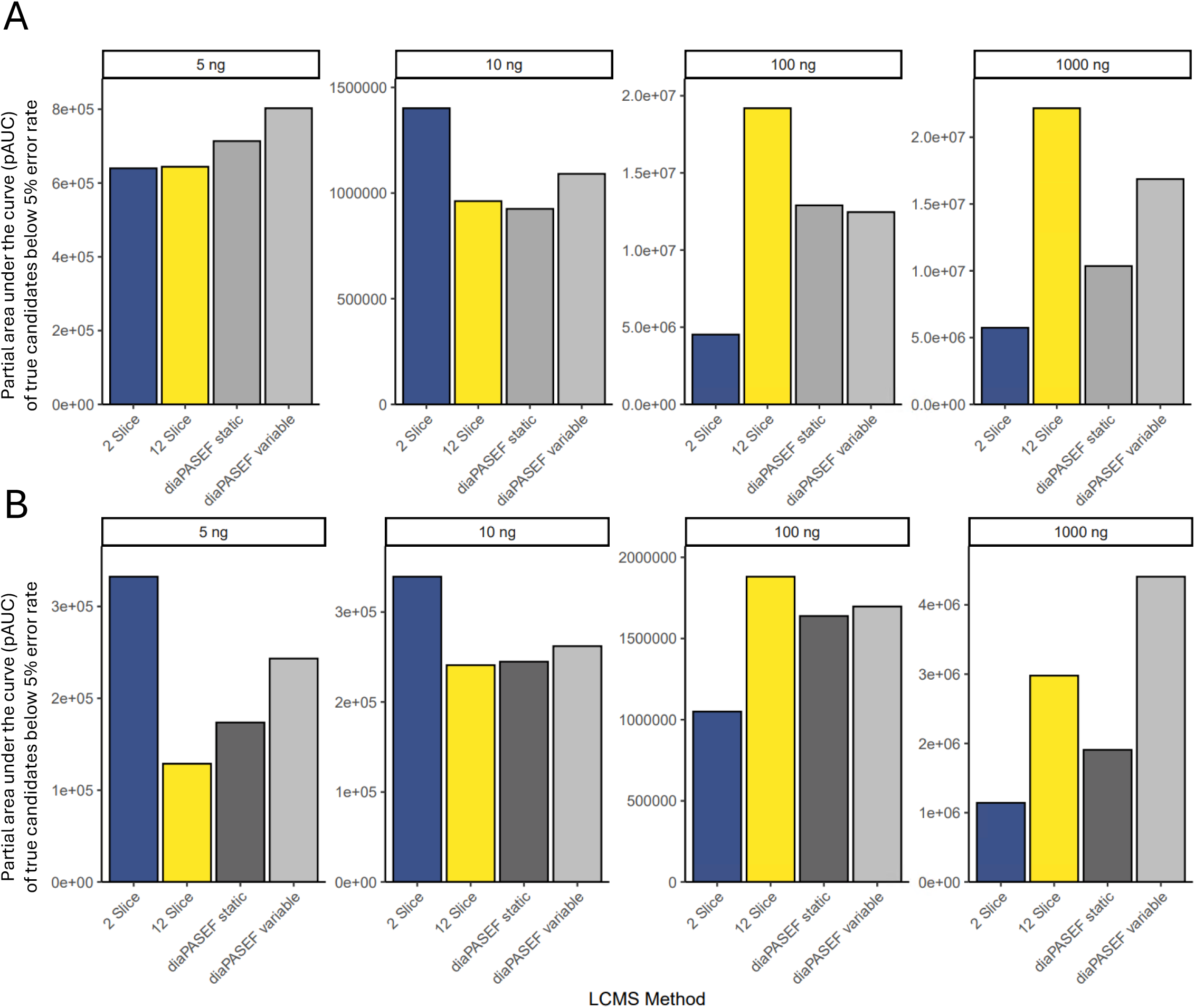
Partial area under the curve of true positive vs candidate length trajectory up to 5% error rate cut-off for selected diagonal-PASEF and dia-PASEF methods at all tested loadings on peptide (**A**) or protein group level (**B**).

